# CENP-C-Mis12 complex establishes a regulatory loop through Aurora B for chromosome segregation

**DOI:** 10.1101/2024.05.29.596395

**Authors:** Weixia Kong, Masatoshi Hara, Yurika Tokunaga, Kazuhiro Okumura, Yasuhiro Hirano, Jiahang Miao, Yusuke Takenoshita, Masakazu Hashimoto, Hiroshi Sasaki, Toshihiko Fujimori, Yuichi Wakabayashi, Tatsuo Fukagawa

**Author notes:** Correspondence to: Masatoshi Hara or Tatsuo Fukagawa.

## Abstract

Establishing the correct kinetochore-microtubule attachment is crucial for faithful chromosome segregation. The kinetochore has various regulatory mechanisms for establishing correct bipolar attachment. However, how the regulations are coupled is not fully understood. Here, we demonstrate a regulatory loop between the kinetochore protein CENP-C and Aurora B kinase, which is critical for the error correction of kinetochore-microtubule attachment. This regulatory loop is mediated through the binding of CENP-C to the outer kinetochore Mis12 complex (Mis12C). Although the Mis12C binding region of CENP-C is dispensable for mouse development and proliferation in human RPE-1 cells, those cells lacking this region display increased mitotic defects. The CENP-C-Mis12C interaction facilitates the centromeric recruitment of Aurora B and the mitotic error correction in human cells. Given that Aurora B reinforces the CENP-C-Mis12C interaction, our findings reveal a positive regulatory loop between Aurora B recruitment and the CENP-C-Mis12C interaction, which ensures chromosome bi-orientation for accurate chromosome segregation.

## Introduction

Accurate chromosome segregation during mitosis is crucial for the transmission of genomic information to the daughter cells. Failure of this process causes chromosomal instability, leading to aneuploidy, a hallmark of cancer (Baker et al., 2024; Bakhoum and Cantley, 2018; Ben-David and Amon, 2020; Girish et al., 2023; Vishwakarma and McManus, 2020).

The kinetochore assembled on the centromere is a large protein complex that establishes a bi-oriented chromosome-microtubule attachment for accurate chromosome segregation during mitosis and meiosis. The kinetochore is composed of two major complexes: the constitutive centromere-associated network (CCAN) at the inner kinetochore and KNL1, Mis12, and Ndc80 complexes (KMN network, KMN for short) at the outer kinetochore (Fukagawa and Earnshaw, 2014; Hara and Fukagawa, 2017; Hara and Fukagawa, 2018; McKinley and Cheeseman, 2016; Mellone and Fachinetti, 2021). The vertebrate CCAN, a 16-protein complex, associates with the centromeric chromatin containing the histone H3 variant CENP-A and establishes a base of the kinetochore structure (Amano et al., 2009; Black and Cleveland, 2011; Foltz et al., 2006; Hori et al., 2008; Izuta et al., 2006; Nishino et al., 2012; Okada et al., 2006; Palmer et al., 1987; Westhorpe and Straight, 2013). During the late G2 and M phases, KMN, the major microtubule-binding complex, is recruited onto CCAN. Consequently, the fully assembled mitotic kinetochore connects the chromosomes with spindle microtubules for chromosome segregation (Alushin et al., 2010; Cheeseman et al., 2006; DeLuca et al., 2006; Hara and Fukagawa, 2017; McKinley and Cheeseman, 2016; Nagpal and Fukagawa, 2016; Pesenti et al., 2016).

The CCAN protein CENP-C is a conserved essential protein for chromosome segregation (Earnshaw and Rothfield, 1985; Fukagawa and Brown, 1997; Fukagawa et al., 1999; Kalitsis et al., 1998; Meluh and Koshland, 1995; Tomkiel et al., 1994) and is thought to be a central hub of kinetochore assembly because CENP-C binds to proteins in all layers of the kinetochore, including CENP-A, other CCAN proteins, and the Mis12 complex (Mis12C) of KMN through its multiple functional domains (Kato et al., 2013; Klare et al., 2015; Screpanti et al., 2011). The interaction between CENP-C and Mis12C was reconstituted in vitro, and structural analyses identified key residues of CENP-C for Mis12-binding, which are conserved among species (Dimitrova et al., 2016; Petrovic et al., 2016). The CENP-C-Mis12C interaction is enhanced by Aurora B-mediated phosphorylation of DSN1, a component of Mis12C, and this phospho-regulation is conserved between yeasts and vertebrates (Bonner et al., 2019; Dimitrova et al., 2016; Hara et al., 2018; Kim and Yu, 2015; Petrovic et al., 2016; Rago et al., 2015). Mis12C also interacts with the microtubule-binding Ndc80 complex (Ndc80C) (Cheeseman et al., 2006; Kline et al., 2006). Therefore, the CENP-C-Mis12C interaction appears to be crucial for bridging centromeres with microtubules. However, this was not the case in chicken DT40 cells. We previously showed that deletion of the Mis12C-binding domain (M12BD) of CENP-C resulted in no apparent growth defects in chicken DT40 cells (Hara et al., 2018). Instead, deletion of the KMN-binding domain from CENP-T, another CCAN protein, causes severe mitotic arrest and consequent cell death, indicating that KMN binding of CENP-T is essential for bioriented chromosome-microtubule attachment in DT40 cells (Hara et al., 2018).

However, given that the key residues in the M12BD of CENP-C and Aurora B-mediated regulation are well conserved among species, unrevealed advantages to the CENP-C-Mis12C interaction that could not be detected in our previous systems may exist. Clarifying the benefits of the CENP-C-Mis12C interaction and elucidating its physiological role is crucial for understanding kinetochore regulation leading to faithful chromosome segregation.

To address these issues, we generated and characterized mice lacking the M12BD of CENP-C (*Cenpc^ΔM12BD^*). Although the M12BD was largely dispensable for mouse development, the *Cenpc^ΔM12BD/ΔM12BD^*mice were caner-prone in the two-stage skin carcinogenesis model, suggesting the contribution of the CENP-C-Mis12C interaction to cancer prevention. This is in line with increased mitotic defects in MEFs established from *Cenpc^ΔM12BD/ΔM12BD^* embryos. To further investigate its molecular mechanisms, we utilized human RPE-1 cells and found that deletion of M12BD reduced the centromeric localization of Aurora B during mitosis. Given that Aurora B kinase regulates kinetochore-microtubule interactions, M12BD deletion resulted in impaired error correction of kinetochore-microtubule attachment, suggesting that the CENP-C-Mis12C interaction positively regulates mitotic Aurora B localization to establish chromosome biorientation. We further clarified the regulatory axis for Aurora B localization using HeLa cells, which are cancerous cells with chromosomal instability and low Aurora B kinase activity at the mitotic centromeres. Forced binding of Mis12C to CENP-C increased Aurora B levels at centromeres and improved error correction efficiency in HeLa cells. Given that Aurora B facilitates the CENP-C-Mis12C interaction (Dimitrova et al., 2016; Kim and Yu, 2015; Petrovic et al., 2016; Rago et al., 2015), we propose a positive regulatory loop between the CENP-C-Mis12C interaction and Aurora B recruitment at the centromeres to maintain Aurora B kinase activity for efficient mitotic error correction to establish bipolar attachment and subsequent faithful chromosome segregation during mitosis.

## Results

### Mis12C-binding domain of CENP-C is dispensable for mouse development but is required for proper mitotic progression in mouse embryonic fibroblasts (MEFs)

CENP-C binds to Mis12C via its N-terminal region (Dimitrova et al., 2016; Hara et al., 2018; Petrovic et al., 2016; Przewloka et al., 2011; Screpanti et al., 2011) (Mis12C-binding domain: M12BD; Figure 1A and Supplementary Figure 1A). We previously found that M12BD is dispensable for the proliferation of chicken DT40 cells (Hara et al., 2018). However, given the amino acid (aa) sequence conservation of M12BD among species and its importance in KMN interactions (Figure 1A and Supplementary Figure 1A), we wondered whether this finding was specific to DT40 cells.

**Figure 1.**
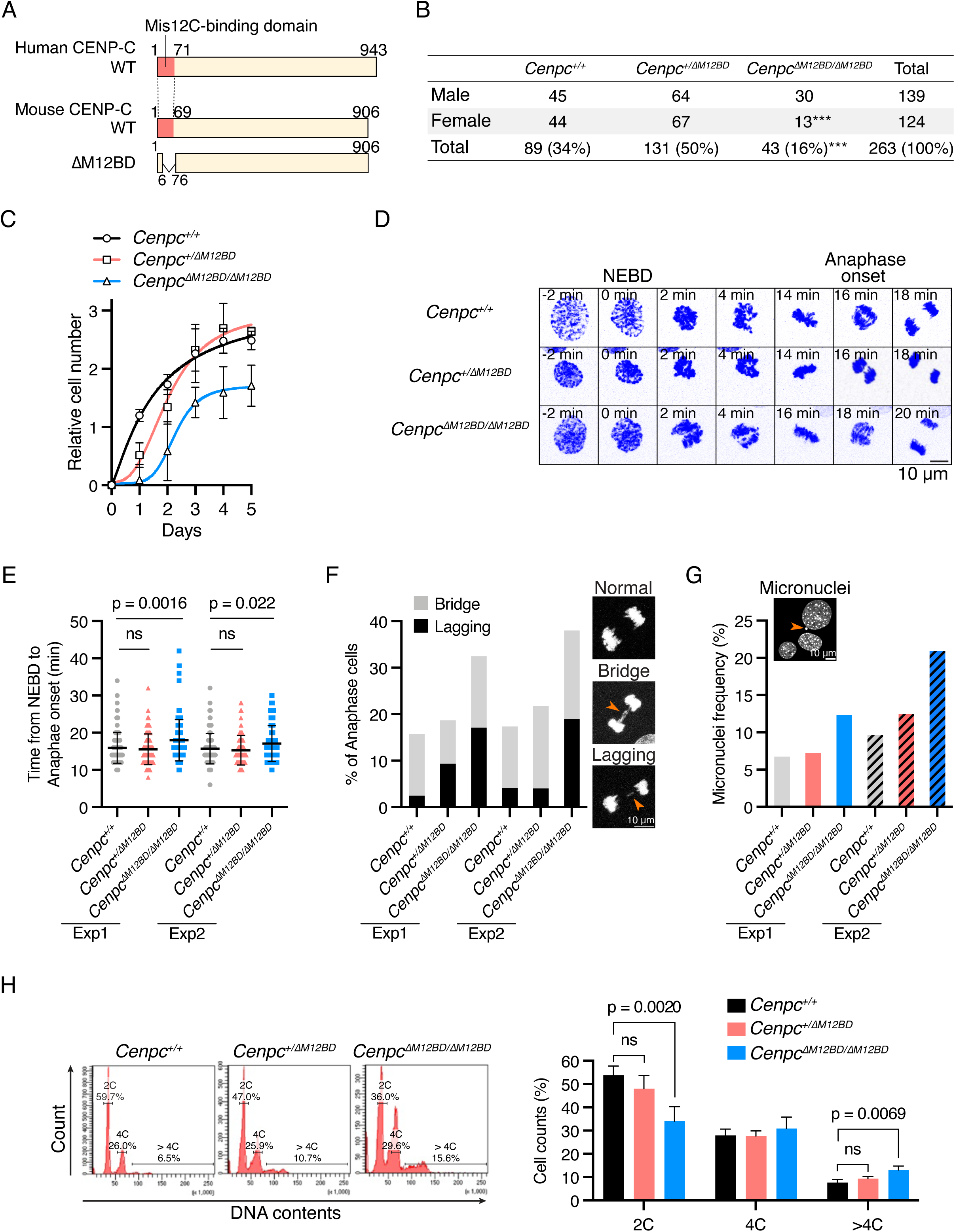
Mis12C-binding domain of CENP-C is dispensable for mouse development but is required for proper mitotic progression in MEFs. (A) Schematic representation of human and mouse CENP-C proteins. The Mis12-binding domain (M12BD) of human CENP-C and its homologous region in mouse CENP-C are highlighted in each CENP-C wild-type (WT). To establish *Cenpc1* (*Cenpc*) mutant mice lacking M12BD (*Cenpc^ΔM12BD^*), using CRISPR/Cas9 genome editing, exon2-4 encoding amino acids 7-75 were deleted from the *Cenpc* gene locus (CENP-C ΔM12BD, see also Supplementary Figure 1). (B) Genotype of offspring from *Cenpc^+/ΔM12BD^* intercross (Chi-squared test. ***, p < 0.001). (C) The growth curve of the MEFs isolated from *Cenpc^+/+^*, *Cenpc^+/ΔM12BD^*, or *Cenpc^ΔM12BD/ΔM12BD^* embryos. The cell numbers were normalized to those at Day 0 of each line. Error bars indicate the mean and standard deviation (SD). (D) Representative time-lapse images of mitotic progression in *Cenpc^+/+^*, *Cenpc^+/ΔM12BD^*, or *Cenpc^ΔM12BD/ΔM12BD^* MEFs. DNA was visualized by staining with SPY650-DNA. Images were projected using maximum intensity projection and deconvoluted. Time is relative to nuclear envelope breakdown (NEBD). Scale bar, 10 μm. (E) Mitotic duration from NEBD to anaphase onset in *Cenpc^+/+^*, *Cenpc^+/ΔM12BD^*, or *Cenpc^ΔM12BD/ΔM12BD^* MEFs. The time-lapse images were analyzed to measure the time from NEBD to anaphase onset. Two independent experiments were performed (Mean and SD, one-way ANOVA with Dunnett’s multiple comparison test, exp1: n = 121 (*Cenpc^+/+^*), 107 (*Cenpc^+/ΔM12BD^*), 117 (*Cenpc^ΔM12BD/ΔM12BD^*), exp2: n = 121 (*Cenpc^+/+^*), 124 (*Cenpc^+/ΔM12BD^*), 121 (*Cenpc^ΔM12BD/ΔM12BD^*)). (F) Chromosome segregation errors in *Cenpc^+/+^*, *Cenpc^+/ΔM12BD^*, or *Cenpc^ΔM12BD/ΔM12BD^* MEFs. The lagging chromosomes and chromosome bridges during anaphase in the cells analyzed in (E) were scored. Representative images are shown. Scale bar, 10 μm. (G) Micronuclei formation in *Cenpc^+/+^*, *Cenpc^+/ΔM12BD^*, or *Cenpc^ΔM12BD/ΔM12BD^* MEFs. MEFs were fixed and the interphase cells with micronuclei were scored (exp1: n = 1570 (*Cenpc^+/+^*), 1753 (*Cenpc^+/ΔM12BD^*), 1679 (*Cenpc^ΔM12BD/ΔM12BD^*), exp2: n = 818 (*Cenpc^+/+^*), 754 (*Cenpc^+/ΔM12BD^*), 765 (*Cenpc^ΔM12BD/ΔM12BD^*)). Representative images are shown. Scale bar, 10 μm. (H) Flow cytometry analysis of DNA contents in *Cenpc^+/+^*, *Cenpc^+/ΔM12BD^*, or *Cenpc^ΔM12BD/ΔM12BD^* MEFs. MEFs from exp1 were fixed and stained with propidium iodide and analyzed by flow cytometry. Representative results are shown on the left. Each cell line was tested in triplicate and analyzed (Mean and SD, one-way ANOVA with Dunnett’s multiple comparison test).

To test whether CENP-C requires M12BD for its functions in other species and to determine the physiological importance of the CENP-C-Mis12C interaction, we generated a mutant mouse model lacking the M12BD of CENP-C. Using the CRISPR/Cas9 system, we deleted exons 2–4 from *Cenpc1* (*Cenpc*) gene, which encodes aa 7-75 (Supplementary Figure 1B). Their deletion removed most of the M12BD, including the key conserved residues for Mis12C-binding (CENP-C^ΔM12BD^; Supplementary Figure 1A and 1B) (Petrovic et al., 2016; Screpanti et al., 2011). In contrast to *Cenpc* null mice, which do not survive embryonic development (Kalitsis et al., 1998), intercrossing heterozygous (*Cenpc^+/11M12BD^*) mice produced homozygous progeny (*Cenpc^11M12BD/11M12BD^*), despite a slight reduction in female offspring (Figure 1B and Supplementary Figure 1C). These results suggest that M12BD of CENP-C is largely dispensable in mouse development; however, female embryos are sensitive to the loss of M12BD.

Chromosomal instability and subsequent micronuclei formation cause female-biased lethality in mouse embryos (McNairn et al., 2019). This prompted us to investigate mitotic progression in MEFs established from E14.5 *Cenpc^11M12BD/11M12BD^* embryos (Supplementary Figure 1D and 1E). We also prepared MEFs from *Cenpc^+/+^* and *Cenpc^+/11M12BD^* mice and confirmed their genotypes and CENP-C protein expression (Supplementary Figure 1E).

We examined the growth of MEFs and found that the proliferation of *Cenpc^11M12BD/11M12BD^* MEFs was slower than that of *Cenpc^+/+^* or *Cenpc^+/11M12BD^* MEFs (Figure 1C). Next, we examined the mitotic progression of MEFs using time-lapse imaging (Figure 1D). It took approximately 16 min from nuclear envelope breakdown (NEBD) to anaphase onset in *Cenpc^+/+^* and *Cenpc^+/11M12BD^*MEFs. The mitotic progression was slightly, but significantly, delayed in *Cenpc^11M12BD/11M12BD^* MEFs (Figure 1D and 1E). We also observed an increase in the population with chromosome mis-segregation during anaphase (lagging chromosome or chromosome bridge) in *Cenpc^11M12BD/11M12BD^* MEFs and in the population with micronuclei in *Cenpc^11M12BD/11M12BD^* MEFs (Figure 1F and 1G). Consistent with these observations, the population with more than 4C DNA content increased in *Cenpc^11M12BD/11M12BD^* MEFs (Figure 1H). These results showed that *Cenpc^11M12BD/11M12BD^* MEFs exhibited chromosomal instability, suggesting that the M12BD of CENP-C contributes to accurate chromosome segregation in mouse cells despite being largely dispensable for development.

### Deletion of M12BD of CENP-C accelerates tumor formation and malignant conversion in the two-stage skin carcinogenesis model

The chromosomal instability is a hallmark of cancer (Baker et al., 2024; Bakhoum and Cantley, 2018; Ben-David and Amon, 2020; Girish et al., 2023; Vishwakarma and McManus, 2020). Since the deletion of M12BD from CENP-C led to increased chromosomal instability in MEFs, we examined the cancer susceptibility of *Cenpc^11M12BD/11M12BD^* mice using a two-stage skin carcinogenesis model (Figure 2A) (Abel et al., 2009; Kemp, 2005). Between 14 and 20 weeks after the initial 7,12-dimethylbenz(a)anthracene (DMBA)/12-O-tetradecanoylphorbol-13-acetate (TPA) treatment *Cenpc^11M12BD/11M12BD^* mice formed significantly more papillomas than *Cenpc^+/+^* or *Cenpc^+/11M12BD^*mice (Figure 2B and 2C). We also monitored mice after 20 weeks and found that malignant conversion was significantly promoted in *Cenpc^11M12BD/11M12BD^* mice by 36 weeks compared to that in *Cenpc^+/+^* or *Cenpc^+/11M12BD^*mice (Figure 2D). These results demonstrate that *Cenpc^11M12BD/11M12BD^* mice are cancer-prone, suggesting that M12BD of CENP-C contributes to cancer prevention.

**Figure 2.**
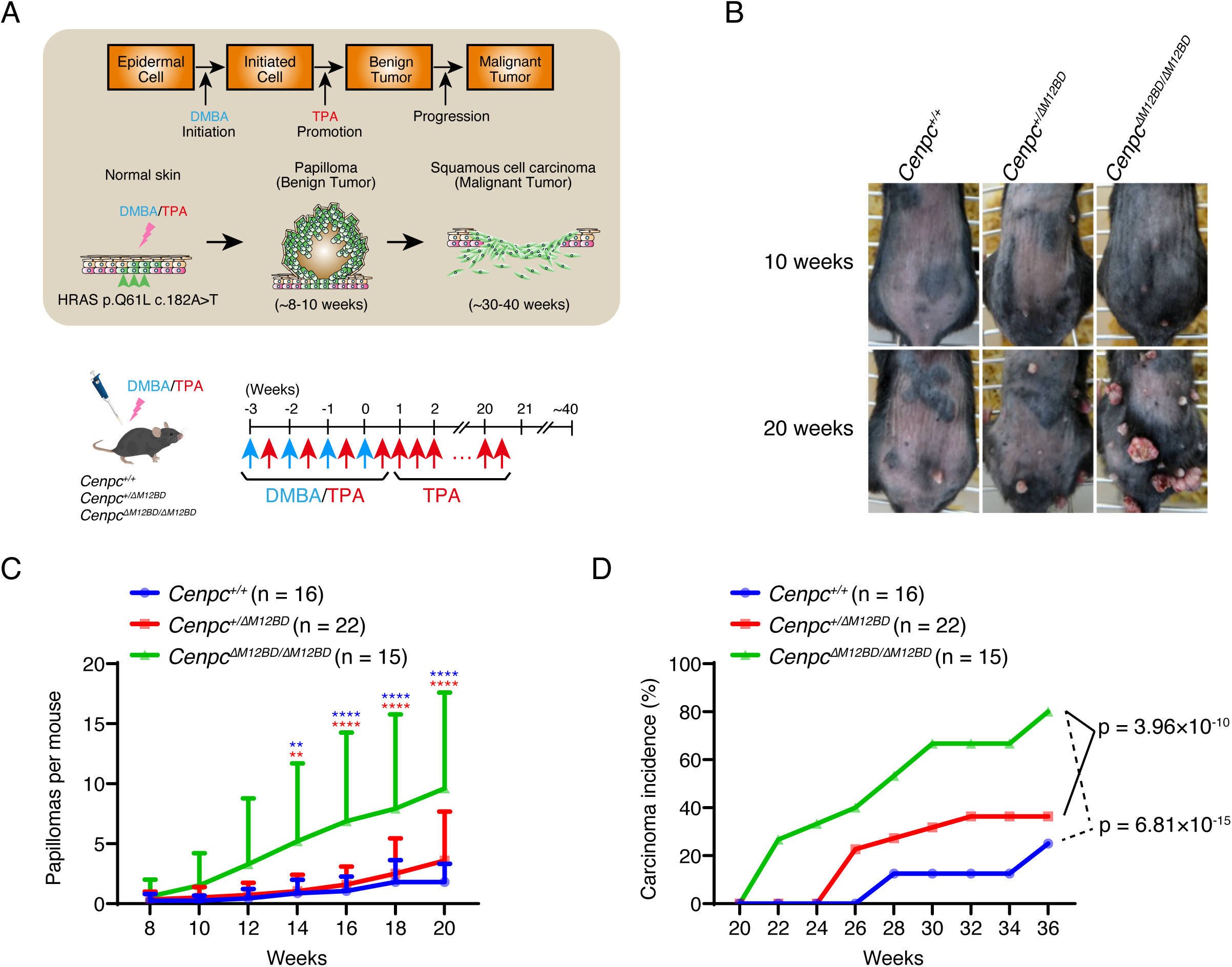
Deletion of the Mis12C-binding domain of CENP-C exacerbates tumor formation and malignant conversion in two-stage skin carcinogenesis model. (A) Schematic representation of the two-stage skin carcinogenesis model. 7,12-dimethylbenz(a)anthracene (DMBA) and 12-O-tetradecanoylphorbol-13-acetate (TPA) were applied to the shaved dorsal back skin. After four rounds of DMBA/TPA treatment, the mice were further treated with TPA to promote papilloma formation and malignant conversion. The pipette and mouse images are adapted from Togo TV (https://togotv.dbcls.jp/en). (B) Representative images of papillomas on the back skin of *Cenpc^+/+^*, *Cenpc^+/ΔM12BD^*, or *Cenpc^ΔM12BD/ΔM12BD^*mice at 10 and 20 weeks after the DMBA/TPA cycles. (C) Papilloma development in *Cenpc^+/+^*, *Cenpc^+/ΔM12BD^*, or *Cenpc^ΔM12BD/ΔM12BD^*mice. The numbers of papillomas on the back skin were counted for each mouse (Mean and SD, two-way ANOVA with Tukey’s test, **: p < 0.01, ****: p < 0.0001, Blue: *Cenpc^+/+^* vs *Cenpc^ΔM12BD/ΔM12BD^*, Red: *Cenpc^+/ΔM12BD^* vs *Cenpc^ΔM12BD/ΔM12BD^*). (D) Malignant conversion in *Cenpc^+/+^*, *Cenpc^+/ΔM12BD^*, or *Cenpc^ΔM12BD/ΔM12BD^* mice. Mice with squamous cell carcinoma were scored from 20 to 36 weeks after the DMBA/TPA cycles (Fisher’s exact test, *Cenpc^+/+^*: n = 16, *Cenpc^+/ΔM12BD^*: n = 22, *Cenpc^ΔM12BD/ΔM12BD^*: n = 15).

### M12BD deletion from CENP-C causes chromosome mis-segregation, leading to mitotic defects in human RPE-1 cells

We aimed to understand the molecular mechanisms by which CENP-C M12BD prevents chromosomal instability. Since MEFs are heterogeneous and their cell proliferation is sensitive to replicative senescence, which limits detailed analyses of mitotic regulation, we used human retinal epithelial cells (RPE-1) for further investigation. RPE-1 is a widely used noncancerous cell line with a stable near-diploid karyotype. We generated RPE-1 cells in which endogenous CENP-C was replaced with FLAG-human CENP-C lacking the Mis12C-binding domain (aa 1-75 region; M12BD; Figure 3A). These cells were referred to as CENP-C^11M12BD^ cells (Figure 3A and Supplementary Figure 2A-G). We also generated RPE-1 cells expressing wild-type FLAG-human CENP-C (CENP-C^WT^ cells) as controls (Figure 3A and Supplementary Figure 2A-G).

**Figure 3.**
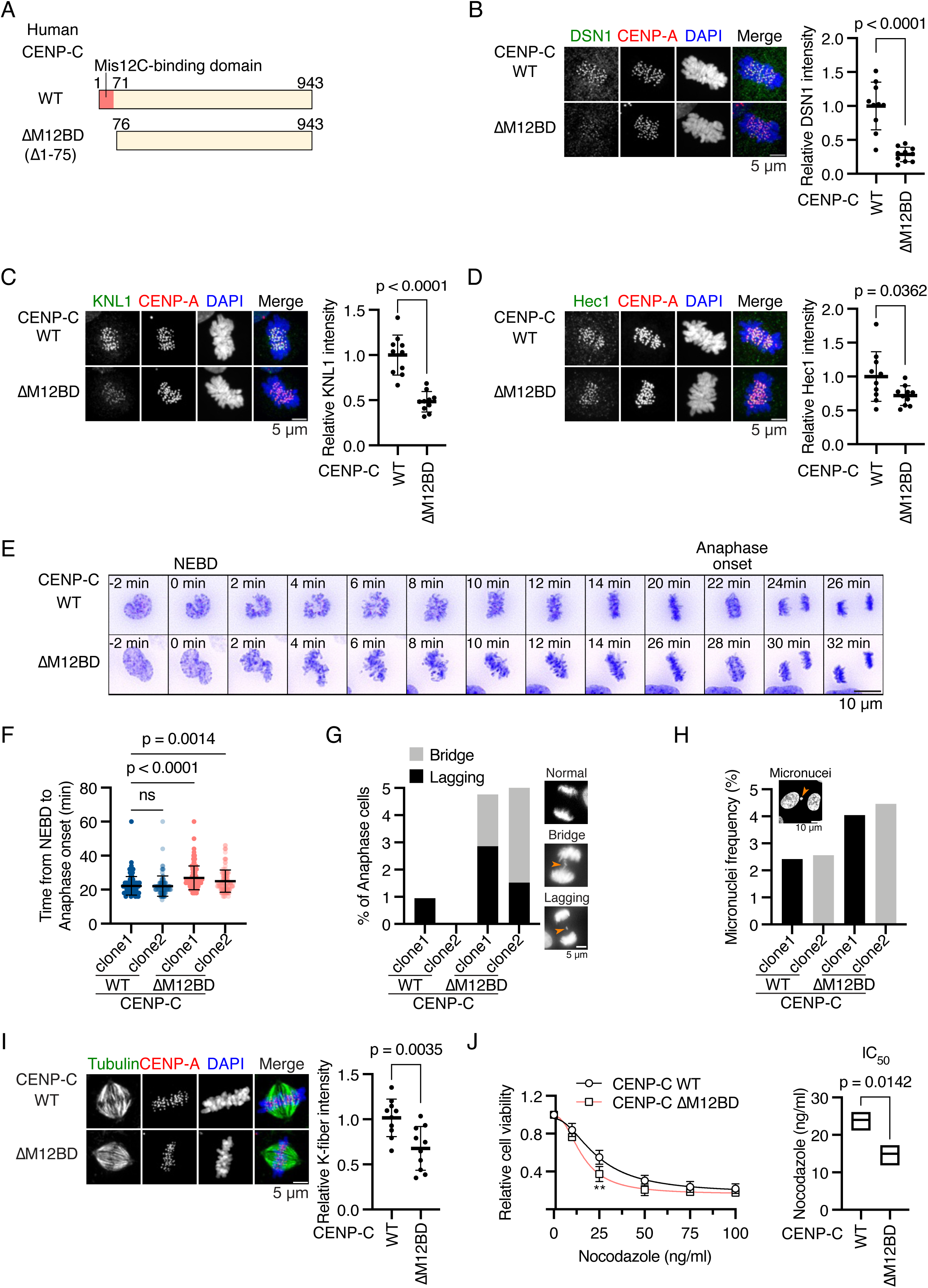
Deletion of the Mis12C-binding domain of human CENP-C delays mitotic progression and increases chromosome segregation errors in RPE-1 cells. (A) Schematic representation of human CENP-C. The Mis12C-binding domain (M12BD, amino acids 1-71) is highlighted in CENP-C wild-type (WT). The N-terminus region (amino acid 1-75) was deleted in CENP-C^ΔM12BD^. FLAG-tagged CENP-C WT or ΔM12BD was introduced into the *CENP-C* locus in RPE-1 cells expressing mScarlet-CENP-A and GFP-H2A (CENP-C^WT^ or CENP-C^ΔM12BD^ cells, respectively. See Supplementary Figure 2). (B) DSN1 localization in CENP-C^WT^ or CENP-C^ΔM12BD^ cells. DSN1, a subunit of the Mis12 complex, was stained with an antibody against DSN1 (green). mScarlet-CENP-A is a kinetochore marker (CENP-A, red). DNA was stained with DAPI (blue). Scale bar, 5 μm. DSN1 signal intensities at mitotic kinetochores were quantified (Mean and SD, two-tailed Student’s t-test, CENP-C^WT^ cells: n = 10, CENP-C^ΔM12BD^ cells: n = 10). (C) KNL1 localization in CENP-C^WT^ or CENP-C^ΔM12BD^ cells. KNL1, a subunit of the KNL1 complex, was stained with an antibody against KNL1. KNL1 localization at mitotic kinetochores was examined and quantified as in (B). Scale bar, 5 μm. Mean and SD, two-tailed Student’s t-test, CENP-C^WT^ cells: n = 10, CENP-C^ΔM12BD^ cells: n = 10. (D) Hec1 localization in CENP-C^WT^ or CENP-C^ΔM12BD^ cells. Hec1, a subunit of the Ndc80C, was stained with an antibody against Hec1. Hec1 localization at mitotic kinetochores was examined and quantified as in (B). Scale bar, 5 μm. Mean and SD, two-tailed Student’s t-test, CENP-C^WT^ cells: n = 10, CENP-C^ΔM12BD^ cells: n = 10. (E) Representative time-lapse images of mitotic progression in CENP-C^WT^ or CENP-C^ΔM12BD^ cells. DNA was visualized with GFP-H2A. Images were projected using maximum intensity projection and deconvoluted. Time is relative to nuclear envelope breakdown (NEBD). Scale bar, 10 μm. (F) Mitotic duration from NEBD to anaphase onset in CENP-C^WT^ or CENP-C^ΔM12BD^ cells. The time-lapse images were analyzed to measure the time from NEBD to anaphase onset. Two independent clones of CENP-C^WT^ or CENP-C^ΔM12BD^ cells were tested (Mean and SD, two-tailed Student’s t-test, CENP-C^WT^ cells clone1: n = 120, CENP-C^WT^ cells clone2: n = 105, CENP-C^ΔM12BD^ cells clone1: n = 105, CENP-C^ΔM12BD^ cells clone2: n = 132). (G) Chromosome segregation errors in CENP-C^WT^ or CENP-C^ΔM12BD^ cells. The lagging chromosomes and chromosome bridges during anaphase in the cells analyzed in (F) were scored. Representative images are shown. Scale bar 5 μm. (H) Micronuclei formation CENP-C^WT^ or CENP-C^ΔM12BD^ cells. The cells were fixed and the interphase cells with micronuclei were scored (CENP-C^WT^ cells clone1: n = 1527, CENP-C^WT^ cells clone2: n = 976, CENP-C^ΔM12BD^ cells clone1: n = 1633, CENP-C^ΔM12BD^ cells clone2: n = 1076). Scale bar, 10 μm. (I) K-fiber in CENP-C^WT^ or CENP-C^ΔM12BD^ cells. CENP-C^WT^ or CENP-C^ΔM12BD^ cells expressing mScarlet CENP-A were fixed after CaCl_2_ treatment and stained with an anti-alpha-tubulin antibody. Scale bar, 5 μm. The mean tubulin signal intensities of the spindle in a cell were quantified as K-fiber signals (Mean and SD, two-tailed Student’s t-test, CENP-C^WT^ cells: n = 10, CENP-C^ΔM12BD^ cells: n = 10). (J) Cell viability of CENP-C^WT^ or CENP-C^ΔM12BD^ cells treated with various concentrations of nocodazole. Viable cells were measured three days after nocodazole addition. Three independent experiments were performed (Mean and SD, two-way ANOVA with Šídák’s multiple comparison test, **: p = 0.0022). IC_50_ indicates the average of concentration to reduce cell viability to 50% from three independent experiments (Mean and SD, two-tailed Student’s t-test).

First, we quantified the Mis12C levels in the kinetochores of CENP-C^11M12BD^ cells. For this, we immunostained DSN1, a component of Mis12C, in CENP-C^11M12BD^ or CENP-C^WT^ cells expressing mScarlet-CENP-A as a kinetochore marker (Supplementary Figure 2A). As shown in Figure 3B, the punctate DSN1 signals found in CENP-C^WT^ cells were significantly reduced in CENP-C^11M12BD^ cells. We also examined the levels of KNL1 and Ndc80 complexes (KNL1C and Ndc80C) by immunostaining with antibodies against their components (KNL1 and Hec1, respectively) and found that the signals of both KNL1C and Ndc80C were reduced in CENP-C^11M12BD^ cells (Figure 3C and 3D). The reduction of Ndc80C levels was mild compared with that of Mis12C and KNL1C. This can be explained by three additional Ndc80C binding sites in CENP-T (Huis In ’t Veld et al., 2016; Rago et al., 2015; Takenoshita et al., 2022).

Next, we examined whether CENP-C^11M12BD^ cells showed mitotic defects as observed in *Cenpc^11M12BD/11M12BD^*MEFs. In contrast to MEFs, in which the deletion of M12BD from CENP-C delayed cell growth, CENP-C^11M12BD^ cells grew comparably to CENP-C^WT^ cells (Supplementary Figure 2H). However, time-lapse imaging showed that the mitotic progression from NEBD to anaphase onset was significantly delayed in CENP-C^11M12BD^ cells, as observed in *Cenpc^11M12BD/11M12BD^* MEFs (Figure 3E and 3F). We also observed an increase in chromosome mis-segregation with lagging or bridging chromosomes in CENP-C^11M12BD^ cells (Figure 3G). In addition, the cell population with micronuclei was increased in CENP-C^11M12BD^ cells (Figure 3H).

### Reduced Ndc80C and kinetochores-associated microtubules (K-fiber) do not cause significant mitotic errors

To clarify the cause of chromosome segregation errors and mitotic delay in CENP-C^11M12BD^ cells, we first examined the K-fiber, as the levels of Ndc80C, which is a critical microtubule-binding complex, were significantly reduced in CENP-C^11M12BD^ cells. Following calcium treatment to depolymerize the highly dynamic microtubules, we stained the remaining stable microtubules, which corresponded to K-fiber, and found that the K-fiber signal intensities in CENP-C^11M12BD^ cells were significantly lower than those in CENP-C^WT^ cells, suggesting a reduction in K-fiber in CENP-C^11M12BD^ cells (Figure 3I). The CENP-C^11M12BD^ cells were more sensitive to low-dose nocodazole treatment than the CENP-C^WT^ cells (Figure 3J). This result further supported the reduction of K-fiber in CENP-C^11M12BD^ cells.

Next, we examined whether the K-fiber reduction in CENP-C^11M12BD^ cells caused mitotic defects in these cells. We used CENP-T mutant cells lacking one Ndc80C binding site. We generated CENP-T mutant RPE-1 cell lines, in which full-length human CENP-T fused with auxin-inducible degron (AID)-tag was expressed from the *AAVS1* locus, and *mScarlet*-fused mutant *human CENP-T* cDNAs lacking either one of two Ndc80C-binding sites (NBD-1: aa 6-31, NBD-2: aa 76-105) were introduced into the endogenous *CENP-T* locus (Figure 4A and Supplementary Figure 3A-3E, *CENP-T^11NBD-1^* or *CENP-T^11NBD-2^*). Upon addition of auxin (IAA), AID-tagged CENP-T was degraded, and these cells expressed only the mScarlet-fused CENP-T mutant protein (Supplementary Figure 3F).

**Figure 4.**
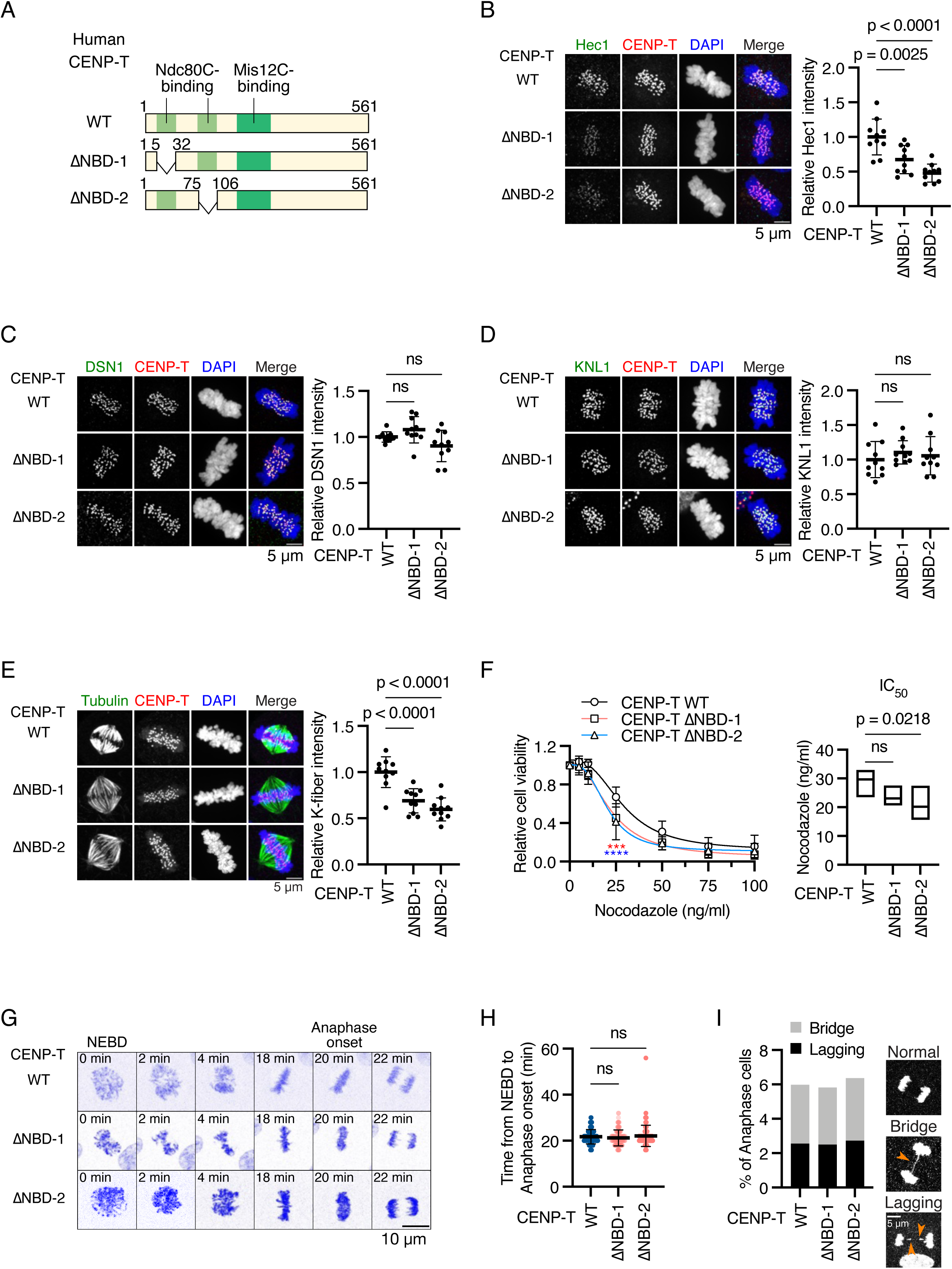
Deletion of one Ndc80C-binding domain of human CENP-T reduces Ndc80C localization and K-fiber but does not cause mitotic defects in RPE-1 cells. (A) Schematic representation of human CENP-T. human CENP-T wild-type (WT) has two Ndc80C binding regions (NBD-1 or -2: amino acids 6-31 or 76-105) and a Mis12-binding domain. Each NBD was deleted in CENP-T^ΔNBD-1^ or CENP-T^ΔNBD-2^, respectively. mScarlet-fused CENP-T WT or each mutant was introduced into *CENP-T* locus in RPE-1 cells expressing OsTIR1 and GFP-mAID-CENP-T (CENP-T^WT^, CENP-T^ΔNBD-1^, or CENP-T^ΔNBD-2^ cells, respectively. See Supplementary Figure 3). (B) Hec1 localization in CENP-T^WT^, CENP-T^ΔNBD-1^, or CENP-T^ΔNBD-2^ cells. Hec1, a subunit of Ndc80C, was stained with an antibody against Hec1 (green). mScarlet-CENP-T is a kinetochore marker (CENP-T, red). DNA was stained with DAPI (blue). Scale bar, 5 μm. Hec1 signal intensities at mitotic kinetochores were quantified (Mean and SD, one-way ANOVA with Dunnett’s multiple comparison test, CENP-T^WT^ cells: n = 10, CENP-T^ΔNBD-1^ cells: n = 10, CENP-T^ΔNBD-2^ cells: n = 10). (C) DSN1 localization in CENP-T^WT^, CENP-T^ΔNBD-1^, or CENP-T^ΔNBD-2^ cells. DSN1, a subunit of Mis12C, was stained with an antibody against DSN1. DSN1 localization at mitotic kinetochores was examined and quantified as in (B). Scale bar, 5 μm. Mean and SD, one-way ANOVA with Dunnett’s multiple comparison test, CENP-T^WT^ cells: n = 10, CENP-T^ΔNBD-1^ cells: n = 10, CENP-T^ΔNBD-2^ cells: n = 10. (D) KNL1 localization in CENP-T^WT^, CENP-T^ΔNBD-1^, or CENP-T^ΔNBD-2^ cells. KNL1, a subunit of KNL1C, was stained with an antibody against KNL1. KNL1 localization at mitotic kinetochores was examined and quantified as in (B). Scale bar, 5 μm. Mean and SD, one-way ANOVA with Dunnett’s multiple comparison test, CENP-T^WT^ cells: n = 10, CENP-T^ΔNBD-1^ cells: n = 10, CENP-T^ΔNBD-2^ cells: n = 10. (E) K-fiber in CENP-T^WT^, CENP-T^ΔNBD-1^, or CENP-T^ΔNBD-2^ cells. CENP-T^WT^, CENP-T^ΔNBD-1^, or CENP-T^ΔNBD-2^ cells were fixed after CaCl_2_ treatment and stained with an anti-alpha-tubulin antibody. CENP-T fused with mScarlet is a kinetochore marker (CENP-T, red). Scale bar, 5 μm. The mean tubulin signal intensities of the spindle in a cell were quantified as K-fiber signals (Mean and SD, one-way ANOVA with Dunnett’s multiple comparison test, CENP-T^WT^ cells: n = 10, CENP-T^ΔNBD-1^ cells: n = 10, CENP-T^ΔNBD-2^ cells: n = 10). (F) Cell viability of CENP-T^WT^, CENP-T^ΔNBD-1^, or CENP-T^ΔNBD-2^ cells treated with various concentrations of nocodazole. Viable cells were measured three days after nocodazole addition. Four independent experiments were performed (Mean and SD, two-way ANOVA with Dunnett’s multiple comparison test, ***: p = 0.0007, ****: p < 0.0001. Red: WT vs ΔNBD-1, Blue: WT vs ΔNBD-2). IC_50_ indicates the average of nocodazole concentration to reduce cell viability to 50% from four independent experiments (Mean and SD, one-way ANOVA with Dunnett’s multiple comparison test). (G) Representative time-lapse images of mitotic progression in CENP-T^WT^, CENP-T^ΔNBD-1^, or CENP-T^ΔNBD-2^ cells. DNA was visualized with SPY505-DNA. Images were projected using maximum intensity projection and deconvoluted. Time is relative to nuclear envelope breakdown (NEBD). Scale bar, 10 μm. (H) Mitotic duration from NEBD to anaphase onset in CENP-T^WT^, CENP-T^ΔNBD-1^, or CENP-T^ΔNBD-2^ cells. The time-lapse images were analyzed to measure the time from NEBD to anaphase onset (Mean and SD, one-way ANOVA with Dunnett’s multiple comparison test, CENP-T^WT^ cells: n = 117, CENP-T^ΔNBD-1^ cells: n = 120, CENP-T^ΔNBD-2^ cells: n = 110). (I) Chromosome segregation errors in CENP-T^WT^, CENP-T^ΔNBD-1,^ or CENP-T^ΔNBD-2^ cells. The lagging chromosomes and chromosome bridges during anaphase in the cells analyzed in (H) were scored. Representative images are shown. Scale bar 5 μm.

In cells expressing CENP-T^11NBD-1^ (CENP-T^11NBD-1^ cells) or CENP-T^11NBD-2^ (CENP-T^11NBD-2^ cells), Ndc80C levels (Hec1) at the kinetochores were significantly lower than those in cells expressing wild-type CENP-T (CENP-T^WT^ cells) (Figure 4B). Importantly, the Mis12C (DSN1) and KNL1C (KNL1) levels at the kinetochores in CENP-T^11NBD-1^ and CENP-T^11NBD-2^ cells were comparable to those in CENP-T^WT^ cells (Figure 4C and 4D). As Ndc80C was reduced at the kinetochores, the K-fiber signal intensities in CENP-T^11NBD-1^ or CENP-T^11NBD-2^ cells were significantly lower than those in CENP-T^WT^ cells. Consistent with K-fiber reduction, CENP-T^11NBD-1^ and CENP-T^11NBD-2^ cells showed sensitivity to low-dose nocodazole (Figure 4F).

Despite the reduced Ndc80C and associated K-fiber reduction, CENP-T^11NBD-1^ and CENP-T^11NBD-2^ cells showed neither significant mitotic delay nor increased chromosome segregation errors (Figure 4G-4I). These results suggest that the K-fiber reduction due to the decrease in Ndc80C is not the cause of mitotic defects in CENP-C^11M12BD^ cells.

### Aurora B localization to mitotic centromeres is diminished in CENP-C^11M12BD^ cells

To determine the cause of chromosome segregation errors and mitotic delay in CENP-C^11M12BD^ cells, we examined the regulatory mechanisms of kinetochore-microtubule attachment. Aurora B is a conserved mitotic kinase that phosphorylates kinetochore substrates, such as Hec1, facilitating the correction of erroneous kinetochore-microtubule attachment (Cheeseman et al., 2006; DeLuca et al., 2006; DeLuca et al., 2011; Liu et al., 2009; Long et al., 2017; Zaytsev et al., 2014). As shown in Figure 5A, Aurora B was localized to the inner centromeric region between sister kinetochores in mitotic cells. We found that Aurora B levels were significantly reduced in CENP-C^11M12BD^ cells compared to those in CENP-C^WT^ cells (Figure 5A). In contrast, the reduction of Aurora B at the centromeres was not observed in CENP-T^11NBD-1^ or CENP-T^11NBD-2^ cells (Figure 5B), which underwent proper mitotic progression despite the reduction of Ndc80C levels at kinetochores (Figure 4).

**Figure 5.**
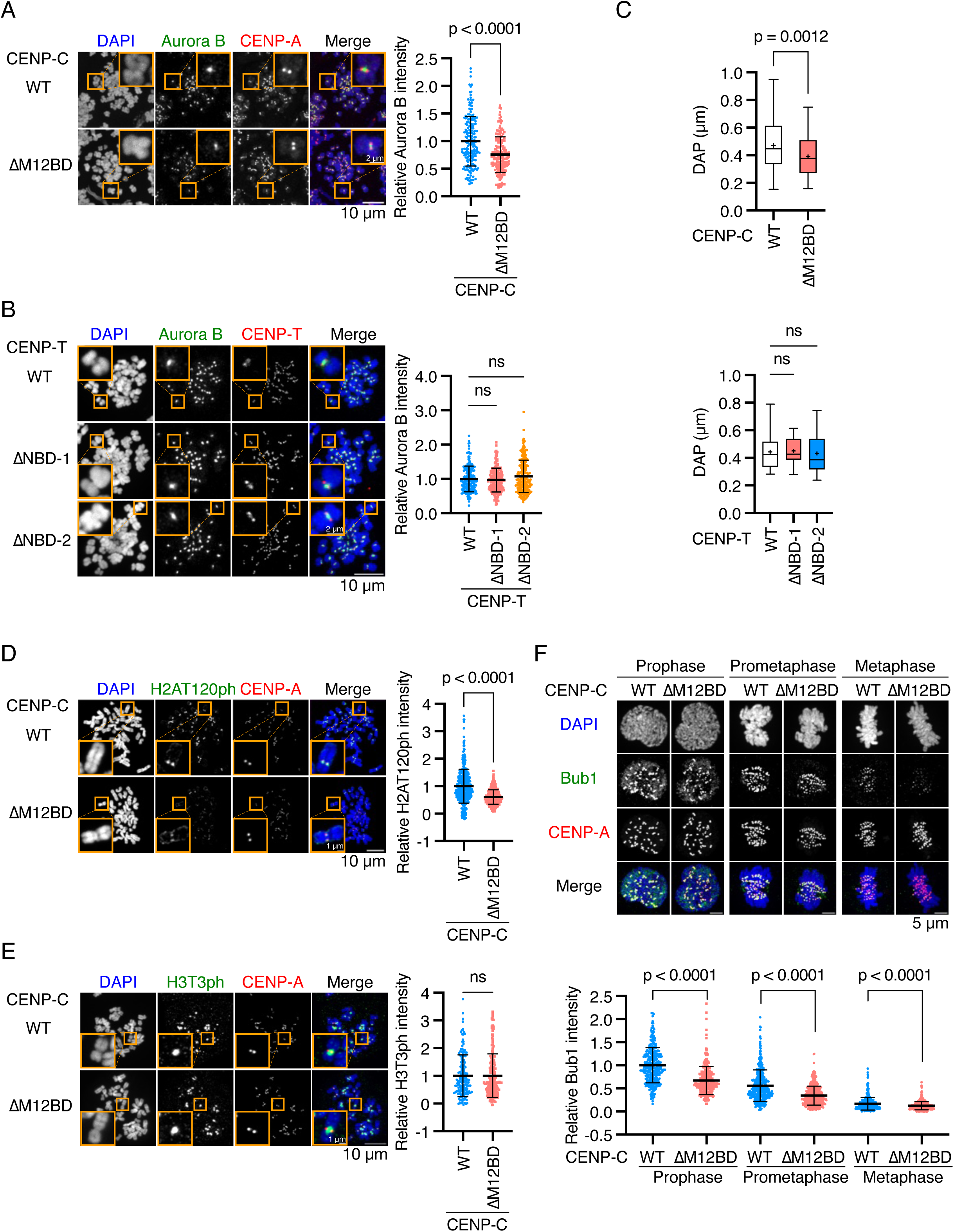
Deletion of Mis12C-binding domain of human CENP-C reduces Aurora B and Bub1 levels at centromeres in RPE-1 cells. (A) Aurora B localization in CENP-C^WT^ or CENP-C^ΔM12BD^ cells. Aurora B was stained with an antibody against Aurora B (green). mScarlet-CENP-A is a kinetochore marker (CENP-A, red). DNA was stained with DAPI (blue). Scale bar, 10 μm. The insets show an enlarged single chromosome (Scale bar, 2 μm). Aurora B signal intensities at inner centromeres were quantified (Mean and SD, two-tailed Student’s t-test, CENP-C^WT^: n = 188 centromeres from 5 cells, CENP-C^ΔM12BD^: n = 166 centromeres from 5 cells). (B) Aurora B localization in CENP-T^WT^, CENP-T^ΔNBD-1^, or CENP-T^ΔNBD-2^ cells. Aurora B and DNA were stained as in (A). mScarlet-CENP-T is a kinetochore marker (CENP-T, red). Scale bar, 5 μm. The Aurora B signal intensities were quantified (Mean and SD, one-way ANOVA with Dunnett’s multiple comparisons test, CENP-T^WT^: n = 197 centromeres from 5 cells, CENP-T^ΔNBD-1^: n = 198 centromeres from 5 cells, CENP-T^ΔNBD-2^: n = 225 centromeres from 5 cells). (C) Chromosome oscillation in CENP-C^WT^, CENP-C^ΔM12BD^, CENP-T^WT^, CENP-T^ΔNBD-1^, or CENP-T^ΔNBD-2^ cells. The deviation from average position (DAP) was calculated from the time-lapse images (Supplementary Figure 4). The graphs display the median and quantile with max and min (+ indicates mean) (Two-tailed Student’s t-test, CENP-C^WT^ cells: n = 54 kinetochore pairs from 12 cells, CENP-C^ΔM12BD^ cells: n = 42 kinetochore pairs from 7 cells; One-way ANOVA with Dunnett’s multiple comparison test, CENP-T^WT^ cells: n = 17 kinetochore pairs from 3 cells, CENP-T^ΔNBD-1^ cells: n = 15 kinetochore pairs from 3 cells, CENP-T^ΔNBD-2^ cells: n = 16 kinetochore pairs from 4 cells). (D) H2AT120ph localization in CENP-C^WT^ or CENP-C^ΔM12BD^ cells. H2AT120ph was stained with an antibody against H2AT120ph (green). mScarlet-CENP-A is a kinetochore marker (CENP-A, red). DNA was stained with DAPI (blue). Scale bar, 10 μm. The insets show an enlarged single chromosome (Scale bar, 1 μm). H2AT120ph signal intensities at kinetochore-proximal centromeres were quantified (Mean and SD, two-tailed Student’s t-test, CENP-C^WT^: n = 398 kinetochores from 5 cells, CENP-C^ΔM12BD^: n = 373 kinetochores from 5 cells). (E) H3T3ph localization in CENP-C^WT^ or CENP-C^ΔM12BD^ cells. H3T3ph was stained with an antibody against H3T3ph. H3T3ph localization at centromeres was examined and quantified as in (D). Scale bar, 10 μm. The graph displayed Mean and SD (Two-tailed Student’s t-test, CENP-C^WT^: n = 170 centromeres from 5 cells, CENP-C^ΔM12BD^: n = 204 centromeres from 5 cells). (F) Bub1 localization in CENP-C^WT^ or CENP-C^ΔM12BD^ cells. Bub1 was stained with an antibody against Bub1 (green). mScarlet-CENP-A is a kinetochore marker (CENP-A, red). DNA was stained with DAPI (blue). Scale bar, 5 μm. Bub1 signal intensities at kinetochores were quantified (Mean and SD, two-tailed Student’s t-test, CENP-C^WT^: n = 318 kinetochores from 5 cells (prophase), n = 354 kinetochores from 5 cells (prometaphase), n = 440 kinetochores from 5 cells (metaphase); CENP-C^ΔM12BD^: n = 280 kinetochores from 5 cells (prophase), n = 423 kinetochores from 5 cells (prometaphase), n = 429 kinetochores from 5 cells (metaphase)).

To further evaluate the reduction of Aurora B activity at the centromeres in CENP-C^11M12BD^ cells, we examined metaphase chromosome oscillation, which is regulated by Aurora B through the phosphorylation of Hec1 (Zaytsev et al., 2014). CENP-C^WT^ or CENP-C^11M12BD^ cells were treated with MG132, a proteasome inhibitor, to arrest the cells at metaphase, and chromosome oscillations were observed by time-lapse imaging (Supplementary Figure 4A). The oscillation amplitude was assessed by quantifying the deviation from the average position (DAP) (Stumpff et al., 2008) for tracked kinetochores labeled with mScarlet-CENP-A (Supplementary Figure 4B and 4C). The amplitude in CENP-C^11M12BD^ was significantly smaller than that in CENP-C^WT^ cells (Figure 5C). In contrast, CENP-T^NBD-1^ and CENP-T^NBD-2^ cells did not show changes in the amplitude of oscillations compared to CENP-T^WT^ cells (Figure 5C and Supplementary Figure 4D). These results suggest a reduction in Aurora B activity at the centromeres in CENP-C^11M12BD^ cells.

Centromeric localization of Aurora B is promoted by the phosphorylation of histone H3 at threonine 3 (H3T3ph) by Haspin and histone H2A at threonine 120 (H2AT120ph) by Bub1 (Wang et al., 2011; Watanabe, 2010). To investigate how the CENP-C-Mis12C interaction is related to Aurora B localization, we examined the phosphorylation of these histones and found that H2AT120ph levels were significantly reduced in CENP-C^11M12BD^ cells; however, H3T3ph levels in CENP-C^11M12BD^ cells were comparable to those in CENP-C^WT^ cells (Figure 5D and 5E). Consistent with the changes in H2AT120ph, CENP-C^11M12BD^ cells showed less Bub1 localization at the kinetochores during mitotic progression than did CENP-C^WT^ cells (Figure 5F). Since Bub1 localizes to kinetochores through KNL1 (Kiyomitsu et al., 2007), the results aligned with the reduction of KNL1C, which binds to Mis12C (Petrovic et al., 2016; Petrovic et al., 2014; Petrovic et al., 2010) at kinetochores in CENP-C^11M12BD^ cells (Figure 3C).

In contrast, Bub1, H2AT120ph, and H3T3ph levels in CENP-T^11NBD-1^ and CENP-T^11NBD-2^ cells, which showed neither mitotic defects nor Aurora B reduction, were comparable to those in CENP-T^WT^ cells (Supplementary Figure 5). The unaltered Bub1 and H2AT120ph levels were consistent with the finding that Mis12C and KNL1C levels at the centromeres in CENP-T^11NBD-1^ and CENP-T^11NBD-2^ cells were comparable to those in CENP-T^WT^ cells (Figure 4C and 4D).

These data suggest that the CENP-C-Mis12C interaction positively regulates centromeric Aurora B localization through the Bub1-H2AT120ph axis.

### Error correction efficiency was reduced in CENP-C^11M12BD^ cells

As one of the key functions of Aurora B is mitotic error correction, which resolves erroneous kinetochore-microtubule attachments to facilitate the correct bipolar spindle microtubule attachment to the sister kinetochores (Cimini et al., 2006; Dewar et al., 2004; Lampson et al., 2004), we evaluated the error correction efficiency in CENP-C^11M12BD^ cells with low Aurora B levels at centromeres (Figure 6A). RPE-1 cells were treated with monastrol, a reversible Eg5 inhibitor (Kapoor et al., 2000), to induce monopolar spindles with erroneous kinetochore-microtubule attachment. After release from monastrol, the cells were arrested at metaphase with MG132, and cells with unaligned chromosomes were scored (Lampson et al., 2004) (Figure 6A). As a control for this error correction assay, we treated CENP-C^WT^ cells with AZD1152, an Aurora B inhibitor, and confirmed an increase in cells with unaligned chromosomes 30 min after release (over 10% and 20% with 40 nM and 200 nM AZD1152, respectively), compared to the cells treated with DMSO (less than 10%) (Figure 6A). This observation is consistent with previous reports, demonstrating that efficient error correction requires Aurora B (Cimini et al., 2006; Lampson et al., 2004).

**Figure 6.**
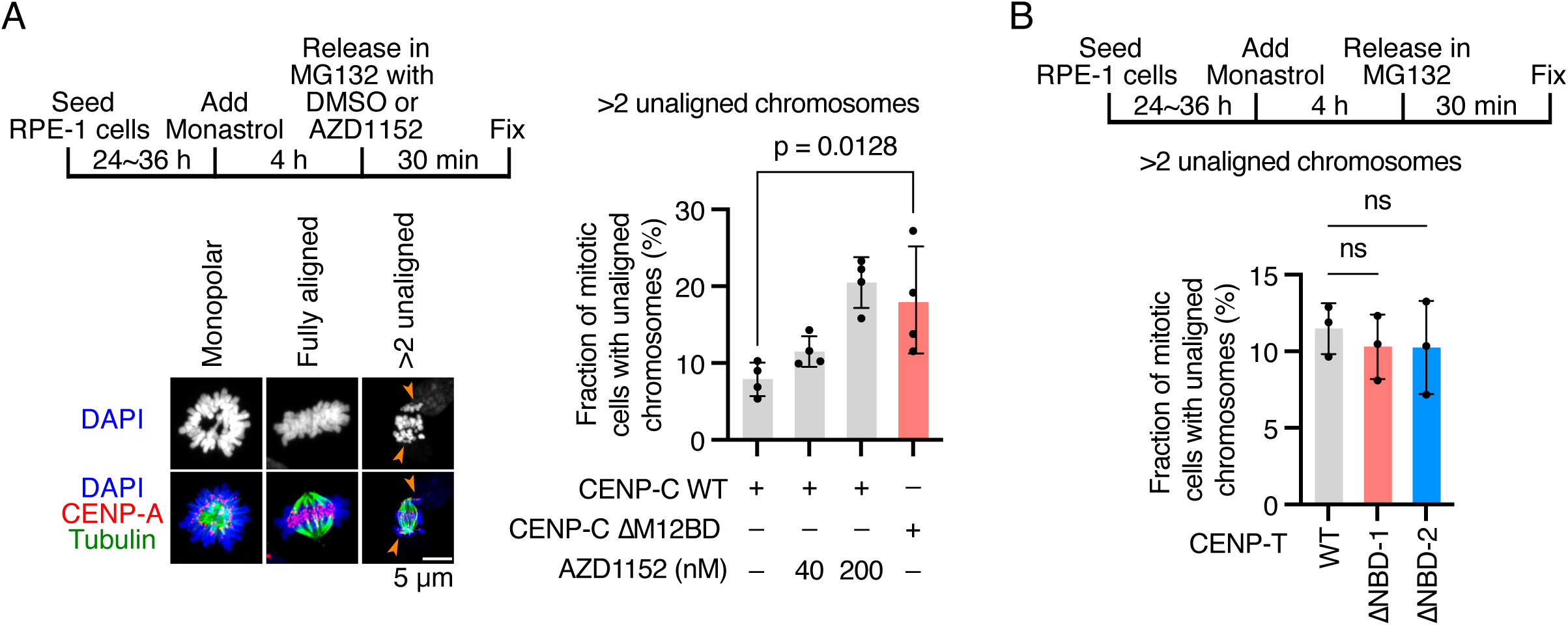
Deletion of Mis12C-binding domain of human CENP-C reduces kinetochore-microtubule error correction efficiency. (A) Error correction assay in CENP-C^WT^ or CENP-C^ΔM12BD^ cells. The cells were treated with monastrol for 4 h and then released and incubated in a medium with MG132 in the presence or absence of an Aurora B inhibitor (AZD1152 or DMSO) for 30 min. The cells were fixed, and microtubules (green) and DNA (blue) were stained with an antibody against alpha-tubulin and DAPI, respectively. mScarlet-CENP-A is a kinetochore marker (CENP-A, red). The cells with more than two unaligned chromosomes were defined as “cells with unaligned chromosomes.” Representative images are shown: a monastrol-treated cell (monopolar), a cell with fully aligned chromosomes (fully aligned), and a cell with unaligned chromosomes (> 2 unaligned). Arrows indicate unaligned chromosomes. Scale bar 5 μm. Mitotic cells with unaligned chromosomes were quantified. Four independent experiments were performed (Mean and SD, two-tailed Student’s t-test). (B) Error correction assay in CENP-T^WT^, CENP-T^ΔNDB-1^, or CENP-T^ΔNDB-2^ cells. The cells were treated with monastrol, released into a medium with MG132, and unaligned chromosomes were quantified as in (A). Three independent experiments were performed. (Mean and SD, one-way ANOVA with Dunnett’s multiple comparison test).

Next, we examined CENP-C^11M12BD^ cells and found that the cells with unaligned chromosomes were significantly increased to ∼20% 30 min after release from monastrol (Figure 6A), suggesting that error correction was less efficient in CENP-C^11M12BD^ cells and that M12BD plays a part in mitotic error correction to facilitate bipolar attachment. In contrast, CENP-T^11NBD-1^ and CENP-T^11NBD-2^ cells exhibited error correction efficiencies equivalent to those in CENP-T^WT^ cells (Figure 6B). This is consistent with the fact that the deletion of either NBD-1 or -2 did not alter the localization of centromeric Aurora B (Figure 5B). In addition, the results implied that reduced K-fiber and Ndc80C levels did not affect error correction efficiency in this assay.

These results suggest that deletion of the M12BD of CENP-C diminishes error correction, possibly due to the reduction of Aurora B levels at centromeres, which is the primary cause of mitotic delay and chromosome segregation errors in CENP-C^11M12BD^ RPE-1 cells.

### Forced binding of Mis12C to CENP-C suppresses chromosomal instability in HeLa cells

The aforementioned results led to a model in which the CENP-C-Mis12C interaction positively regulates Aurora B localization at centromeres, facilitating the biorientation of chromosomes. To further support this model, we used HeLa cells, a cancer cell line with chromosomal instability and low Aurora B activity at mitotic centromeres (Figure 7A) (Abe et al., 2016). We hypothesized that forcing Mis12C-binding to CENP-C would increase Aurora B levels at the centromeres and ameliorate error correction efficiency in HeLa cells (Figure 7A). To do this, we utilized DSN1 lacking the basic motif, which bypasses the Aurora B-mediated regulation of the CENP-C-Mis12C interaction (Dimitrova et al., 2016; Hara et al., 2018; Kim and Yu, 2015; Petrovic et al., 2016; Rago et al., 2015). The basic motif of DSN1 masks the Mis12C surface for CENP-C binding, and Aurora B phosphorylation of the basic motif releases it from the Mis12C surface to facilitate the CENP-C-Mis12C interaction. Deletion of the basic motif of DSN1 reinforces the CENP-C-Mis12C interaction (Figure 7A) (Hara et al., 2018; Kim and Yu, 2015; Petrovic et al., 2016).

**Figure 7.**
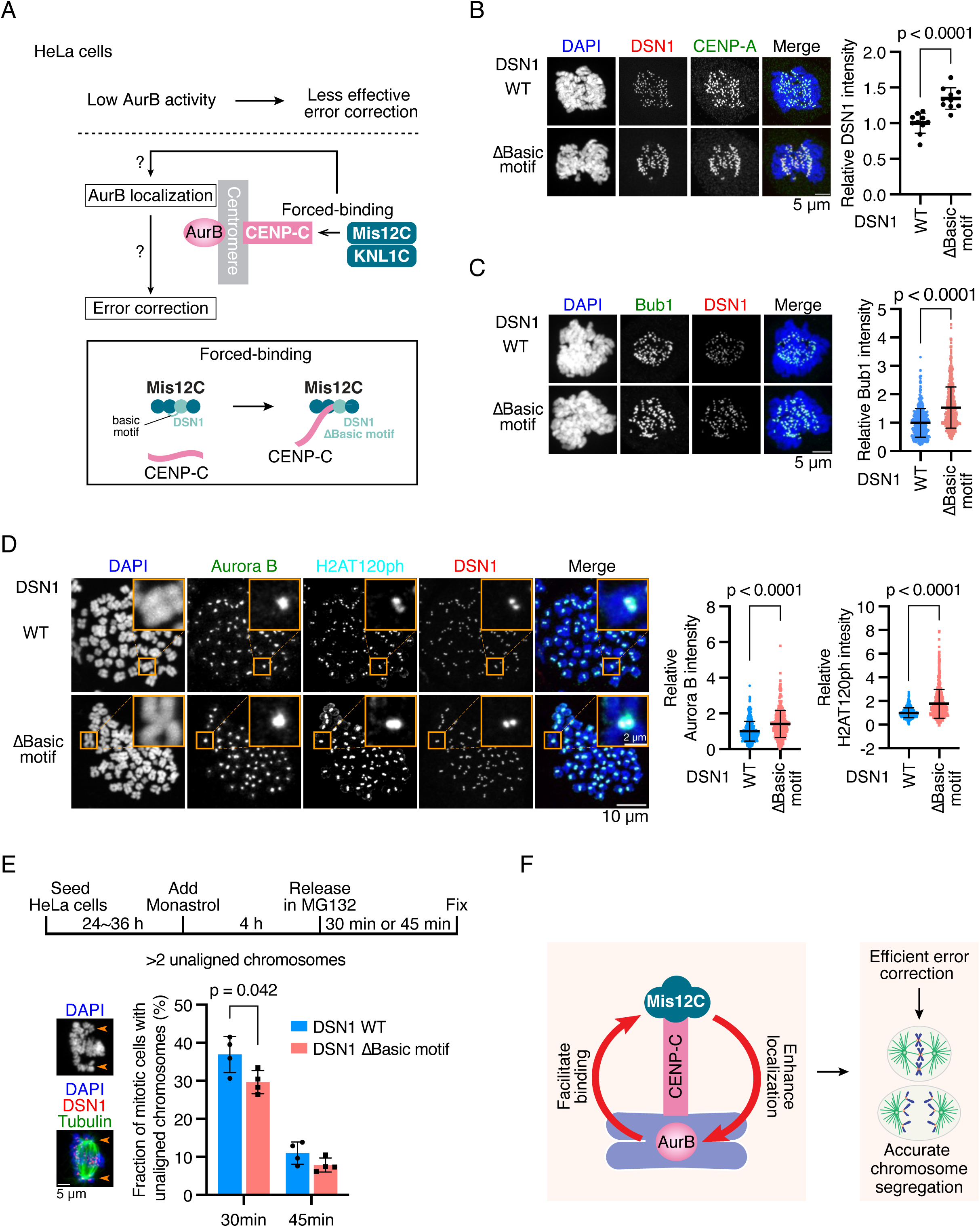
Forced binding of Mis12C to CENP-C increases Aurora B levels on centromeres and improves error correction in HeLa cells. (A) Schematic representation for forced binding of Mis12C to CENP-C in HeLa cells. To validate the idea that the CENP-C-Mis12C interaction positively regulates Aurora B localization, we utilized HeLa cells in which Aurora B activity is low at centromeres, leading to chromosome instability. The CENP-C-Mis12C interaction was increased by expressing a DSN1 mutant lacking the basic motif (ΔBasic motif) in HeLa cells and examined the Aurora B levels and efficiency of kinetochore-microtubule error correction. Dsn1 is a subunit of Mis12C. The basic motif of DSN1 masks the CENP-C-binding surface of Mis12C, preventing the CENP-C-Mis12C interaction. The deletion of the basic motif increases the binding affinity between CENP-C and Mis12C (Petrovic et al., 2016). (B) DSN1 localization in DSN1^WT^ or DSN1^ΔBasic^ ^motif^ HeLa cells. DSN1, a subunit of the Mis12 complex, was stained with an antibody against DSN1 (red). DNA was stained with DAPI (blue). CENP-A was stained by an antibody against CENP-A as a kinetochore marker (green). Scale bar, 5 μm. DSN1 signal intensities at mitotic kinetochores were quantified (Mean and SD, two-tailed Student’s t-test, DSN1-C^WT^: n = 10 cells, DSN1^ΔBasic^ ^motif^: n = 10 cells). (C) Bub1 localization in DSN1^WT^ or DSN1^ΔBasic^ ^motif^ HeLa cells. Bub1 was stained with an antibody against Bub1 (green). DSN1 was stained as a kinetochore marker (red). DNA was stained with DAPI (blue). Scale bar, 5 μm. Bub1 signal intensities at kinetochores were quantified (Mean and SD, two-tailed Student’s t-test, DSN1-C^WT^: n = 523 kinetochores from 6 cells, DSN1^ΔBasic^ ^motif^: n = 501 kinetochores from 6 cells). (D) Aurora B and H2AT120ph localization in DSN1^WT^ or DSN1^ΔBasic^ ^motif^ HeLa cells. Aurora B and H2AT120ph were stained with their antibodies (green and cyan). mScarlet-DSN1 is a kinetochore marker (DSN1, red). DNA was stained with DAPI (blue). Scale bar, 10 μm. The insets show an enlarged single chromosome (Scale bar, 2 μm). The signal intensities of Aurora B at centromeres and H2AT120ph at kinetochore-proximal centromeres were quantified (Mean and SD, two-tailed Student’s t-test, DSN1-C^WT^: n = 385 centromeres, 651 kinetochores from 6 cells for Aurora B, H2AT120ph, respectively, DSN1^ΔBasic^ ^motif^: n = 379 centromeres, 668 kinetochores from 6 cells for Aurora B, H2AT120ph, respectively). (E) Error correction assay in DSN1^WT^ or DSN1^ΔBasic^ ^motif^ HeLa cells. The cells were treated with monastrol for 4 h and then released and incubated in a medium with MG132 for 30 or 45 min. The cells were fixed, and microtubules (green) and DNA (blue) were stained with an antibody against alpha-tubulin and DAPI, respectively. mScarlet-DSN1 is a kinetochore marker (DSN1, red). The cells with more than two unaligned chromosomes were defined as “cells with unaligned chromosomes.” Arrows indicate unaligned chromosomes. Scale bar 5 μm. Mitotic cells with unaligned chromosomes were quantified. Four independent experiments were performed (Mean and SD, two-tailed Student’s t-test). (F) Model for a positive regulatory loop to facilitate Aurora B localization at the centromere through the CENP-C-Mis12C interaction. The regulatory system is required for efficient error correction of kinetochore-microtubule attachment, leading to accurate chromosome segregation.

We generated a HeLa cell line expressing a DSN1 mutant lacking the basic motif (referred to as the DSN1^11Basic^ ^motif^) from the endogenous *DSN1* locus (DSN1^11Basic^ ^motif^ cells; Supplementary Figure 6A-6C). Mis12C (DSN1) levels at kinetochores, as well as KNL1C (KNL1) and Ndc80C (Hec1) levels, were increased in DSN1^11Basic^ ^motif^ cells compared with those in DSN1^WT^ cells (Figure 7B and Supplementary Figure 6D), indicating that a stable CENP-C-Mis12C interaction occurred in Dsn1^11Basic^ ^motif^ cells. This forced binding of Mis12C to CENP-C significantly increased the centromeric localization of Aurora B, Bub1, and H2AT120ph in DSN1^11Basic^ ^motif^ cells (Figure 7C and 7D). Finally, we assessed the error correction efficiency in DSN1^11Basic^ ^motif^ cells and found that it was improved in DSN1^11Basic^ ^motif^ HeLa cells (Figure 7E). These results indicate that forced binding of Mis12C to CENP-C facilitates Aurora B localization at the centromeres and improves error correction efficiency in HeLa cells, supporting the hypothesis that the CENPC-Mis12C interaction positively regulates Aurora B localization at the centromeres.

## Discussion

In this study, we demonstrated that the CENP-C-Mis12C interaction positively regulates Aurora B localization and facilitates mitotic error correction to establish bi-oriented chromosomes. Given that Aurora B regulates CENP-C-Mis12C binding during mitosis through DSN1 phosphorylation, we propose a feedback mechanism to control Aurora B levels at the centromeres during mitosis (Figure 7F). Communication between the inner centromere and outer kinetochore through CENP-C is crucial for maintaining sufficient levels of Aurora B for bi-oriented chromosome establishment and accurate chromosome segregation to prevent chromosomal instability and cancer formation (Figure 7F).

The CENP-C-Mis12C interaction has been studied as a possible crucial platform for recruiting Ndc80C for microtubule binding, and its regulatory mechanisms have been revealed at the molecular level (Dimitrova et al., 2016; Petrovic et al., 2016). Contrary to its proposed importance, *Cenpc^11M12BD/11M12BD^* mice are viable, suggesting that the CENP-C-Mis12C interaction is largely dispensable for mouse development. This is likely because CENP-T is the major scaffold for KMN recruitment to kinetochores in mammalian cells, as in chicken DT40 cells (Hara et al., 2018).

Nevertheless, *Cenpc^11M12BD/11M12BD^* mice are cancer-prone in a two-stage skin carcinogenesis model. This observation suggests that the CENP-C-Mis12C interaction is of physiological importance for ensuring accurate chromosome segregation in cells exposed to stress. We demonstrated that MEFs established from *Cenpc^11M12BD/11M12BD^* embryos showed significant chromosome segregation errors. Our culture conditions were not optimal for MEFs (Parrinello et al., 2003), resulting in high basal chromosome segregation errors even in control cells. We also found that the CENP-C-Mis12C interaction was unnecessary for cell proliferation in RPE-1 cells, a near-diploid human cell line; however, it was required for efficient bi-oriented chromosome establishment.

We previously reported that the CENP-C-Mis12C interaction is dispensable for chicken DT40 cell proliferation (Hara et al., 2018). In contrast to mammalian cells, we did not observe any obvious mitotic defects in DT40 cells lacking M12BD. This may be attributed to technical limitations in detecting subtle differences in chromosome segregation in DT40 cells, which have many tiny chromosomes in small cell volumes. An alternative explanation may be the difference in CENP-C characteristics between mammals and chickens. In human cells, CENP-C depletion causes CCAN disassembly (McKinley et al., 2015), whereas CCAN proteins, including CENP-T, remain on the centromeres in CENP-C-knockout chicken DT40 cells (Hori et al., 2008), implying that other kinetochore proteins compensate CENP-C functions for kinetochore assembly in chicken DT40 cells.

Error correction of kinetochore-microtubule attachment was less efficient in CENP-C^11M12BD^ cells. As deletion of the M12BD of CENP-C reduced Ndc80C levels, as well as Mis12C and KNL1C, Ndc80C recruitment on CENP-C via Mis12C might be crucial for efficient error correction. However, RPE-1 cells expressing CENP-T lacking the Ndc80C binding region showed no defects in error correction or chromosome segregation, indicating that Ndc80C reduction is not a major cause of mitotic defects in RPE-1 CENP-C^11M12BD^ cells. We previously demonstrated that kinetochores contain excess Ndc80C molecules via CENP-T (Takenoshita et al., 2022). The remaining Ndc80C levels in the CENP-C^11M12BD^ cells may be sufficient for efficient error correction. Instead, the CENP-C-Mis12C interaction functions through KNL1C, which recruits Bub1. In CENP-C^ΔM12BD^ cells, Bub1, H2A120ph, and Aurora B levels were reduced in CENP-C^ΔM12BD^ cells, leading to inefficient error correction of kinetochore-microtubule attachment. Bub1-Aurora B reduction was not observed in the cells expressing CENP-T lacking one of the Ndc80C binding regions. As Bub1 also plays a role in the spindle assembly checkpoint machinery, the CENP-C-Mis12C interaction may also facilitate the spindle assembly checkpoint; however, this was not addressed in the current study.

Since CENP-T also recruits Mis12C and KNL1C to kinetochores, the CENP-T-Mis12C interaction may also be involved in the feedback loop that maintains Aurora B (Figure 7F). Although this possibility cannot be ruled out by our current results, we suggest that the Aurora B feedback loop is specifically mediated by the CENP-C-Mis12C interaction. Firstly, the copy number of CENP-C at the kinetochore was higher than that of CENP-T in human cells (CENP-C 215 vs. CENP-T 72) (Suzuki et al., 2015). Secondly, CENP-C is the major scaffold for recruiting Mis12C during prometaphase, during which error correction is performed (Hara et al., 2018). Thirdly, the CENP-C-Mis12C interaction is largely dependent on DSN1 phosphorylation by Aurora B (Kim and Yu, 2015; Petrovic et al., 2016; Rago et al., 2015), whereas the CENP-T-Mis12C interaction is regulated by CDK1-mediated CENP-T phosphorylation and Aurora B-mediated DSN1 phosphorylation (Walstein et al., 2021). CDK1-mediated CENP-T phosphorylation promotes CENP-T-Mis12C interaction without DSN1 phosphorylation by Aurora B in vitro (Huis In ’t Veld et al., 2016; Walstein et al., 2021), suggesting that the CENP-T-Mis12C interaction may be less sensitive to Aurora B activity. Further analyses are necessary to clarify whether CENP-C-Mis12C and CENP-T-Mis12C interactions have distinct roles.

A previous study demonstrated that Aurora B is enriched in the inner centromeres of misaligned chromosomes and reduced in aligned chromosomes in RPE-1 cells (Salimian et al., 2011). This study also showed that the levels of Aurora B-mediated DSN1 phosphorylation are increased at kinetochores on unaligned chromosomes with Aurora B enrichment. Based on these observations, feedback control from the kinetochores to the inner centromeres has been proposed for Aurora B levels to sense chromosome bi-orientation (Salimian et al., 2011). In the present study, we clarified the molecular basis for feedback control: at misaligned chromosomes, Aurora B facilitates DSN1 phosphorylation and the subsequent CENP-C-Mis12C interaction, which turns on the feedback loop to promote Aurora B localization, leading to the correction of erroneous microtubule attachment. Once chromosomes align with the bipolar attachment, Aurora B substrates on the kinetochore, including DSN1, are dephosphorylated through spatial separation mechanisms (Liu et al., 2009; Tanaka, 2002), diminishing the CENP-C-Mis12C interaction and suppressing the regulatory loop. This reduces Aurora B at the aligned chromosomes and leads to further dephosphorylation of its substrates, such as Hec1, which stabilizes the kinetochore-microtubule attachment. The feedback loop control for Aurora B localization is established by the CENP-C-Mis12C interaction, together with the spatial Aurora B separation mechanism, and contributes to efficient error correction for chromosome biorientation.

In C33A cells, a carcinoma cell line, DSN1 has a mutation at Ser109, which is the Aurora B phosphorylation site that regulates the CENP-C-Mis12C interaction (Depmap: https://depmap.org/portal/), suggesting that the CENP-C-Mis12C-mediated Aurora B regulatory loop may be impaired in some cancer cells. Indeed, the control of Aurora B levels on unaligned chromosomes is halted in HeLa cells (Salimian et al., 2011), which may be responsible for chromosomal instability in HeLa cells. This is further supported by our finding that the enforced binding of Mis12C to CENP-C increased Aurora B protein levels at the centromeres and improved error correction efficiency. However, its efficiency was lower than that in RPE-1 cells. The maximum activation of Aurora B kinase requires the binding of the heterochromatin protein HP1 at the mitotic centromeres, and this regulatory mechanism is hampered in cancer cells (Abe et al., 2016), suggesting that in addition to a certain amount of Aurora B at the centromeres, HP1-mediated Aurora B activation is also required to control Aurora B activity for efficient error correction.

The mechanisms controlling Aurora B localization through Bub1/H2AT120ph or the CENP-C-Mis12 interaction by Aurora B have been well studied, separately. Our study highlights that these two mechanisms are integrated into a regulatory system to form a feedback loop for bi-oriented kinetochore-microtubule attachment. This implies that other regulatory mechanisms that function on centromeres/kinetochores would combine to form regulatory systems. A future understanding of how the regulatory mechanisms are integrated will shed light on the dynamic regulatory systems of centromeres/kinetochores to ensure accurate chromosome segregation, providing potential targets for cancer therapy.

### Materials and Methods Establishment of mutant mouse lines

We utilized electroporation to introduce Cas9 protein and sgRNAs into 1-cell embryos (Hashimoto et al., 2016) to delete the exon 2 to 4. Embryos were obtained from superovulated B6C3F1 females crossed to B6C3F1 males. Embryos were aligned in the 1-mm gap of a CUY501G1 electrode (Nepa Gene, Ichikawa, Japan) filled with 200 ng/μL of freshly prepared Guide-it TM Recombinant Cas9 (Electroporation-Ready, Cat#632641, Takara Bio, Japan), 100 ng/µl of each sgRNA in Opti-MEM I (Thermo Fisher Scientific). Electroporation was performed at 30 V (3 msec ON + 97 msec OFF) x 7 pluses using CUY21 EDIT electroporator (BEX, Tokyo, Japan). The embryos were then cultured in modified Whitten’s medium (DR01032, PHC Japan) over night at 37°C under 5% of CO_2_. The embryos that reached the 2-cell stage were transferred to the oviducts of pseudopregnant females. Cesarean sections were performed when pregnant females did not deliver naturally, and pups were raised with ICR foster mothers. After genotyping, mice with the deletion were crossed with C57BL/6 to F1 heterozygous mice.

Animal care and experiments were conducted in accordance with the Guidelines of Animal Experiment of the National Institutes of Natural Sciences and the Guide for the Care and Use of Laboratory Animals of the Ministry of Education, Culture, Sports, Science, and Technology of Japan. The experiments employed in this study were approved by the Institutional Animal Care and Use Committee of the National Institutes of Natural Sciences and by the Committee on the Ethics of Animal Experiments of Chiba Cancer Center.

### gRNA synthesis

To delete exons 2 to 4 of the mouse *Cenpc1* (*Cenpc*) gene, we selected gRNA target sequences intron 1 and intron 4 using CRISPick (Doench et al., 2016; Sanson et al., 2018). gRNAs were synthesized as previously described (Hashimoto et al., 2016). We amplified DNA fragments containing T7 promotor, gRNA target sequence (intron 1: CCAACACTATAGCTGACAAG, intro 4: AAACTGATAGAGTACAGTGG), and gRNA scaffold sequence by PCR using primers shown in Table S1 and pX330 (Addgene plasmid #42230) (Cong et al., 2013) as a template. After purification of the PCR products by ethanol precipitation, gRNAs were synthesized from the PCR products using MEGAshortscript T7 Transcription Kit (Thermo Fisher Scientific) and purified by phenol-chloroform-isoamyl alcohol extraction and isopropanol precipitation. gRNAs used in this study were shown in Table S1.

### Establishment of mouse embryonic fibroblasts (MEFs)

Mouse embryonic fibroblasts were isolated from 14.5-day-old embryos of *Cenpc^+/+^*, *Cenpc^+/ΔM12BD^*, and *Cenpc^ΔM12BD/ΔM12BD^*mice. Following the removal of the head and organs, the embryos were rinsed with PBS, minced, and subjected to digestion with trypsin-EDTA (0.05%) (Gibco) for 30 min at 37°C. Trypsin was quenched by adding DMEM with high glucose (Sigma-Aldrich) supplemented with 15% fetal bovine serum (FBS). Each digested embryo was then plated on a 100-mm-diameter dish and incubated in humidified air containing 5% CO_2_ at 37°C. The passage number was documented for each batch of mouse embryonic fibroblasts and the first passage cells (P1) were cryopreserved in freeze cryopreservation medium (BAMBANKER, GC LYMPHOTEC) at -80°C.

### Two-step skin carcinogenesis

Sixteen *Cenpc^+/+^*, 22 *Cenpc^+/ΔM12BD^*, and 15 *Cenpc^ΔM12BD/ΔM12BD^* mice were subjected to the following protocol. At 8-10 weeks of age, the backs of the mice were shaved with an electric clipper. Two days later, 7,12-dimethylbenz(a)anthracene (DMBA) (Sigma), (25 µg per mouse in 200 µl acetone) was applied to the shaved dorsal back skin. Three days after the first DMBA treatment, 12-*O*-tetradecanoylphorbol-13-acetate (TAP) (Calbiochem), (10 µg per mouse in 200 µl acetone) was administered. After four rounds of DMBA/TPA treatment, the mice were further treated with TPA twice a week for 20 weeks. The number of papillomas was recorded from 8 to 20 weeks, and the development of squamous cell carcinoma was monitored for up to 36 weeks after TPA treatment.

### Cell culture

Human hTERT-RPE-1 and HeLa Kyoto cell lines were maintained in a culture medium containing DMEM (Nacalai Tesque) supplemented with 10% fetal bovine serum (Sigma) and Penicillin-Streptomycin (100 μg/ml) (Thermo Fisher) and cultured at 37°C, 5% CO_2_. For degradation of GFP-mAID-CENP-T (RPE-1 cKO-CENP-T) cells were treated with 500 µM of 3-Indole acetic acid (IAA; Wako).

For the analysis of MEFs, the P1 MEFs were thawed and subcultured in a culture medium containing DMEM (Nacalai Tesque) supplemented with 10% FBS (Sigma) and Penicillin-Streptomycin (100 μg/ml) (Thermo Fisher) at 37°C, 5% CO_2_. They were then passed one more time (P2) and used for all downstream experiments.

### Plasmid constructions for cell transfection

To express mScarlet-fused full-length CENP-A under the control of the CENP-A promoter in RPE-1 cells, the sequence of mScarlet-fused CENP-A followed by puromycin (PuroR) or neomycin resistance genes (NeoR) expression cassette driven by the beta-actin (ACTB) promoter was cloned into the pBluescript II SK (pBSK) with 5’ and 3’ homology arm fragments (approximately 1 kb each) surrounding the *CENP-A* start codon (pBSK_mScarlet-CENP-A) using In-Fusion Snap Assembly Master Mix (Takara Bio). To integrate the construct into the endogenous *CENP-A* locus, CRISPR/Cas9-mediated homologous recombination was utilized, employing pX330 (Addgene plasmid #42230) (Cong et al., 2013) containing single-guide RNA (sgRNA) targeting a genomic sequence (GGGCCTCGGGCTTTCGGCTC) around the CENP-A start codon (pX330_sgCENP-A). sgRNA for CENP-A was designed using CRISPOR (Concordet and Haeussler, 2018).

To express GFP-fused full-length CENP-A under the control of CMV promoter in RPE-1 cells, the sequence of GFP-fused CENP-A followed by L-Histidinol resistance genes (HisD) expression cassette driven by the *ACTB* promoter was cloned into the pT2/HB (a gift from Perry Hackett, Addgene plasmid #26557) (pT2/HB_GFP-CENP-A). The GFP-fused CENP-A with HisD expression cassette was integrated into the genome using the Sleeping Beauty transposon system (Mates et al., 2009).

To express GFP-fused histone H2A from the *AAVS1* locus in RPE-1 cells, the sequence of GFP-fused H2A followed by the blasticidin S resistance gene (*BsR*) expression cassette driven by the *ACTB* promoter was cloned into the pBSK with 5’ and 3’ homology arm fragments of *AAVS1* locus (approximately 1 kb each) (pBSK_GFP-H2A) using In-Fusion Snap Assembly Master Mix (Takara Bio). The construct was integrated into the endogenous *AAVS1* locus by CRISPR/Cas9-mediated homologous recombination, employing pX330 (Addgene plasmid #42230) (Cong et al., 2013) containing sgRNA targeting a genomic sequence (ACCCCACAGTGGGGCCACTA) within intron 1 of *PPP1R112C* (pX330_sgAAVS1). sgRNA for *AAVS1* locus was designed using CRISPOR (Concordet and Haeussler, 2018).

To express Flag-tagged full-length human CENP-C (CENP-C^WT^) or ΔMis12C-binding domain (M12BD: aa1-75) under the control of the endogenous *CENP-C* promoter in RPE-1 cells, the cDNA of *Flag-tagged human CENP-C WT* or *ΔM12BD* followed by the zeocin (*ZeoR*) or *HisD* expression cassette driven by the *ACTB* promoter was cloned into the pBSK with 5’ and 3’ homology arm fragments (approximately 1 kb each) surrounding the *CENP-C* start codon (pBSK_FLAG-CENP-C^WT^ or FLAG-CENP-C^ΔM12BD^) using In-Fusion Snap Assembly Master Mix (Takara Bio). The constructs were integrated into the endogenous *CENP-C* locus by CRISPR/Cas9-mediated homologous recombination, employing pX330 (Addgene plasmid #42230)(Cong et al., 2013) containing sgRNA targeting a genomic sequence (GGCCGGAACATGGCTGCGTC) around the *CENP-C* start codon (pX330_sgCENP-C). sgRNA for *CENP-C* was designed using CRISPOR (Concordet and Haeussler, 2018).

To express OsTIR1-T2A-BsR and GFP-mAID-fused human CENP-T simultaneously under control of the CMV promoter, *CENP-T* cDNA was cloned into the pAID1.2-NEGFP (Nishimura and Fukagawa, 2017; Nishimura et al., 2009; Nishimura et al., 2020) (pAID1.2-CMV-NGFP-CENP-T, which includes CMV promoter-OsTIR1-T2A-BsR-IRES2-GFP-mAID-CENP-T). To express OsTIR1-T2A-BsR and GFP-mAID-fused CENP-T simultaneously under the control of the CMV promoter from the *AAVS1* locus in RPE-1 cells, the sequence of CMV promoter OsTIR1-T2A-BsR-IRES2-GFP-mAID-CENP-T was cloned into the pBSK with 5’ and 3’ homology arm fragments of *AAVS1* locus (approximately 1 kb each) (pBSK_AAVS1_OsTIR1_GFP-mAID-CENP-T). The construct was integrated into the endogenous *AAVS1* locus in RPE-1 cells by CRISPR/Cas9-mediated homologous recombination by using pX330_sgAAVS1.

Mutant CENP-T cDNAs (ΔNBD-1 (Ndc80C-binding domain 1: aa6-31), ΔNBD-2 (Ndc80C-binding domain 2: aa76-105), CENP-T^ΔNBD-1^ and CENP-T^ΔNBD-2^, respectively) were generated using PCR and In-Fusion Snap Assembly Master Mix (Takara Bio). To express the mScarlet-fused full-length CENP-T (CENP-T^WT^), CENP-T^ΔNBD-1^, or CENP-T^ΔNBD-2^ under the control of the endogenous *CENP-T* promoter in RPE-1 cells, the cDNA of *mScarlet-*fused *CENP-T WT*, *ΔNBD-1*, or *ΔNBD-2* followed by *NeoR* or *PuroR* expression cassette driven by the *ACTB* promoter was cloned into the pBSK with 5’ and 3’ homology arm fragments (approximately 1 kb each) surrounding the *CENP-T* start codon (pBSK_mScarlet-CENP-T^WT^, CENP-T^ΔNBD-1^, or CENP-T^ΔNBD-2^). Each construct was integrated into the endogenous *CENP-T* locus in RPE-1 cells by CRISPR/Cas9-mediated homologous recombination, employing pX330 (Addgene plasmid #42230) (Cong et al., 2013) containing sgRNA targeting a genomic sequence (AGACGATGGCTGACCACAAC) around the *CENP-T* start codon (pX330_sgCENP-T). sgRNA for *CENP-T* was designed using CRISPOR (Concordet and Haeussler, 2018).

To express mScarlet-fused full-length DSN1 (DSN1^WT^) or ΔBasic motif (aa91-113) mutant under the control of the endogenous *DSN1* promoter in HeLa cells, the cDNA of *mScarlet*-fused *DSN1 WT* or *ΔBasic motif* followed by *NeoR* and *PuroR* expression cassette driven by the *ACTB* promoter was cloned into the pBSK with 5’ and 3’ homology arm fragments (approximately 1 kb each) surrounding the *DSN1* start codon (pBSK_mScarlet-DSN1^WT^ or DSN1^ΔBasic^ ^motif^) using In-Fusion Snap Assembly Master Mix (Takara Bio). Each construct was integrated into the endogenous *DSN1* locus in HeLa cells by CRISPR/Cas9-mediated homologous recombination, employing pX330 (Addgene plasmid #42230) containing sgRNA targeting a genomic sequence (CTTACCTTGGGTTCAGGCTT) around the *DSN1* start codon (pX330_sgDSN1). sgRNA for DSN1 was designed using CRISPOR (Concordet and Haeussler, 2018).

To express mouse CENP-C protein (aa1-405) in *E. coli*, *mouse CENP-C* (aa1-405) cDNA was cloned into pET30b (Merck) or pGEX6p-1 (Cytiva).

### Generation of cell lines

To establish RPE-1 cell lines expressing mScarlet-fused CENP-A under the control of the endogenous *CENP-A* promoter, the RPE-1 cells were co-transfected with pBSK_mScarlet-CENP-A and pX330_sgCENP-A using Neon Transfection System (Thermo Fisher) with 6 pulses (1400 V, 5 msec), as previously described (Takenoshita et al., 2022). Transfected cells were then subjected to selection in a medium containing 2 mg/ml puromycin (Takara Bio) and 500 μg/ml G418 (Sigma) to isolate single-cell clones (RPE-1 mScarlet-CENP-A cells).

To generate RPE-1 mScarlet-CENP-A cells expressing GFP-fused H2A from the *AAVS1* locus under the control of the CMV promoter, RPE-1 mScarlet-CENP-A cells were co-transfected with pBSK_GFP-H2A and pX330_sgAAVS using Neon Transfection System (Thermo Fisher) with 6 pulses (1400 V, 5 msec). Transfected cells were selected in a medium containing 1 mg/ml Blasticidin S hydrochloride (Kaken Pharmaceutical) to isolate single-cell clones (RPE-1 mScarlet-CENP-A GFP-H2A cells).

To establish RPE-1 mScarlet-CENP-A GFP-H2A cells expressing either CENP-C WT or ΔM12BD under the control of the endogenous *CENP-C* promoter, RPE-1 mScarlet-CENP-A GFP-H2A cells were co-transfected with pBSK_FLAG-CENP-C^WT^ or CENP-C^ΔM12BD^ and pX330_sgCENP-C using Neon Transfection System (Thermo Fisher) with 6 pulses (1400 V, 5 msec). Transfected cells were selected in a medium containing 10 ng/ml Zeocin (Invitrogen) and 1.5 mg/ml L-Histidinol dihydrochloride (Sigma) to isolate single-cell clones (RPE-1 mScarlet-CENP-A GFP-H2A Flag-CENP-C^WT^ or CENP-C^ΔM12BD^ cells).

To generate an inducible CENP-T protein degradation system using the AID system in RPE-1 cells, the CENP-T AID system expression cassette (CMV promoter OsTIR1-T2A-BsR-IRES2-GFP-mAID-CENP-T) was integrated into the *AAVS1* locus. RPE-1 cells were co-transfected with pBSK_AAVS1_OsTIR1_GFP-mAID-CENP-T and pX330_sgAAVS1 using Neon Transfection System (Thermo Fisher) with 6 pulses (1400 V, 5 msec). Transfected cells were selected in a medium containing 1 mg/ml Blasticidin S hydrochloride (Kaken Pharmaceutical) to isolate single-cell clones (RPE-1 cKO-CENP-T cells).

To generate RPE-1 cKO-CENP-T cells (GFP-mAID-CENP-T) expressing either CENP-T WT, ΔNBD-1, or ΔNBD-2, RPE-1 cKO-CENP-T cells were co-transfected with pBSK_mScarlet-CENP-T^WT^, CENP-T^ΔNBD-1^, or CENP-T^ΔNBD-2^, and pX330_sgCENP-T using Neon Transfection System (Thermo Fisher) with 6 pulses (1400 V, 5 msec). Transfected cells were selected in a medium containing 2 mg/ml puromycin (Takara Bio) and 500 μg/ml G418 (Sigma) to isolate single-cell clones (RPE-1 cKO-CENP-T mScarlet-CENP-T^WT^, CENP-T^ΔNDB-1^, or CENP-T^ΔNDB-2^ cells). To express GFP-CENP-A in RPE-1 cKO-CENP-T mScarlet-CENP-T^WT^, CENP-T^ΔNBD-1^, or CENP-T^ΔNBD-2^ cells, we utilized the Sleeping Beauty transposon system (Mates et al., 2009) The cells were transfected with pT2/HB_GFP-CENP-A and pCMV(CAT)T7-SB100 (Addgene plasmid #34879) (Mates et al., 2009) selected in a medium containing 1.5 mg/ml L-Histidinol dihydrochloride (Sigma) to isolate single-cell clones (RPE-1 cKO-CENP-T GFP-CENP-A mScarlet-CENP-T^WT^, CENP-T^ΔNBD-1^, or CENP-_TΔNBD-2)._

To establish HeLa cells expressing mScarlet-fused DSN1 WT or ΔBasic motif under the control of the endogenous *DSN1* promoter, HeLa cells were co-transfected with pBSK_mScarlet-DSN1^WT^ or DSN1^ΔBasic^ ^motif^, and pX330_sgDSN1 using Neon Transfection System (Thermo Fisher) with 6 pulses (1400 V, 5 msec). Transfected cells were selected in a medium containing 2 mg/ml puromycin (Takara Bio) and 2 mg/ml G418 (Sigma) to isolate single-cell clones (HeLa mScarlet-DSN1^WT^ or DSN1^ΔBasic^ ^motif^ cells).

### Cell counting

To count the number of RPE-1 cells or MEFs, the culture medium was aspirated and then the cells were washed with PBS. Subsequently, 2.5 g/liter of Trypsin and 1 mmol/liter EDTA solution (Nacalai Tesque) was added and incubated for 3 to 5 minutes at room temperature (RT). Trypsin was quenched by adding the culture medium. The cells were suspended by pipetting several times and then mixed with an equal volume of 0.4 wt/vol% Trypan Blue Solution (Wako). The living cells were counted using Countess II (Thermo Fisher).

### Genotyping

To extract genomic DNA from MEFs, RPE-1 cells, or HeLa cells, the cells were collected after trypsinization, and washed with PBS as described previously (Takenoshita et al., 2022). The collected cells were resuspended in 0.2 mg/ml Proteinase K (Sigma) in PBST (0.1% Tween 20 (Nacalai tesque) in PBS) and then incubated for 90 min at 55°C followed by heating for 15 min at 96°C. The integration of target constructs was confirmed by PCR using primers listed in Table S1.

### Anti-mouse CENP-C antibody generation

Mouse CENP-C aa1-405 fused with 6 × His was expressed in *E. coli* Rosetta2(DE3) transformed with pET30b-mouse CENP-C^1-405^ and affinity-purified. The purified protein was injected into rabbits to raise antisera (Wako). For affinity purification of mouse CENP-C antibody, GST-fused mouse CENP-C aa1-405 was expressed in E. coli Rosetta2(DE3) transformed with pGEX6P-1-mouse CENP-C^1-405^ and affinity-purified. GST-CENP-C^1-405^ was conjugated with CNBr Sepharose 4B (Cytiva) and incubated with the antiserum for 1 h at RT. After a wash with 50 mM Tris-HCl pH7.5, 150 mM NaCl, the antibodies were eluted by 200 mM Glycin-HCl pH2.0, 150 mM NaCl and immediately neutralized with 1/20 vol. of 1 M Tris. The affinity-purified antibodies were concentrated and then buffer-exchanged to 50 mM Tris-HCl pH7.5, 150 mM NaCl with Amicon Ultra-0.5 ml (Merk).

### Immunoblotting

RPE-1 cells, MEFs, or HeLa cells were collected after trypsinization, washed with PBS, and suspended in 1x Laemmli Sample Buffer (LSB: 62.5 mM Tris (Trizma base, Sigma)-HCl, pH6.8, 2% SDS (Nacalai tesque), 10% Glycerol (Nacalai tesque), 50 mM DTT (Nacalai tesque), bromophenol blue (Wako)) to a final concentration of 1×10^4^ cells/µl. The lysate was sonicated and heated for 5 min at 96°C. Subsequently, the lysate was separated by 5-20% SDS-PAGE (SuperSepAce, Wako) and transferred onto a PVDF membrane (Immobilon-P, Merck). After washing in TBST (0.1% Tween 20 in TBS (50 mM Tris-HCl pH7.5, 150 mM NaCl)) for 15 min, the membrane was incubated with primary antibodies overnight at 4°C. Following a 15 min wash with TBST, the membrane was incubated with secondary antibodies for 1 h at RT. After another 15 min wash with TBST, the membrane was incubated with ECL Prime Western Blotting Detection Reagent (Cytiva) for 5 min. The signal was detected and visualized using a ChemiDoc Touch imaging system (Bio-Rad) and the image was processed using Image Lab (Bio-Rad) and Photoshop 2019 (Adobe).

To detect CENP-C signals in RPE-1 CENP-C^WT^ or CENP-C^ΔM12BD^ cell lines, harvested cells were suspended in TMS buffer (20 mM Tris-HCl, pH8.0, 5 mM MgCl_2_, 250 mM sucrose, 0.5% NP-40, 10% glycerol) for 10 min on ice. The cells were spun down at 17,000 x*g* for 15 min at 4°C. The pellet was collected and washed with TKS buffer (20 mM Tris-HCl, pH8.0, 200 mM KCl, 250 mM sucrose, 1% Triton X-100, 10% glycerol, 1 mM DTT). The precipitate was suspended in lysis buffer (50 mM NH_2_PO_4_, 50 mM Na_2_HPO_4_, 0.3 M NaCl, 0.1% NP-40, 1 mM DTT). Following sonication, the lysate was diluted in 2 x LSB and heated for 5 min at 96°C. The proteins were detected as above.

The primary antibodies used in this study were rabbit anti-mouse CENP-C (1:5000), Guinea pig anti-human CENP-C (1:10,000) (Ando et al., 2002), mouse anti-human CENP-A (1:3000) (Ando et al., 2002), rabbit anti-human DSN1 (1:5000) (a gift from Iain Cheeseman, Whitehead Institute, MIT) (Kline et al., 2006), rat anti-RFP (1:1000) (Chromotek), rat anti-human CENP-T (1:1000) (a gift from Kinya Yoda, Nagoya University, Nagoya, Japan), rat anti-histone H3 (1:5000) (a gift from Hiroshi Kimura, Tokyo Tech, Tokyo, Japan) (Nozawa et al., 2010), and mouse anti-α-tubulin (1:10,000) (Sigma). The secondary antibodies were HRP-conjugated anti-rabbit IgG (1:10,000) (Jackson ImmunoResearch), HRP-conjugated anti-guinea pig IgG (1:10,000) (Sigma), HRP-conjugated anti-mouse IgG (1:10,000) (Jackson ImmunoResearch), and HRP-conjugated anti-rat IgG (1:10,000) (Jackson ImmunoResearch). All antibodies were diluted in Signal Enhancer Hikari (Nacalai tesque) to enhance signal sensitivity and specificity.

### Immunofluorescence staining and image acquisition

For the analysis of DSN1, KNL1, CENP-A, and Bub1, RPE-1 or HeLa cells were seeded onto 35 mm glass-bottom culture dishes (MatTek) for 24 to 36 hours. The samples were fixed with 4% paraformaldehyde (PFA; Electron Microscopy Sciences) in PHEM buffer (60 mM PIPES, 25 mM HEPES, 10 mM EGTA, 2 mM MgCl2, pH6.8) for 10 min at RT. The cells were permeabilized with 0.5% Triton X-100 in PBS for 10 min at RT, followed by blocking with antibody dilution buffer (Abdil, 3% BSA, 0.1% Triton X-100, 0.1% NaAzide in TBS) for 5 to 10 min at RT. The samples were then incubated with primary antibodies for 1 h at 37°C or overnight at 4°C. After washing the samples with PBST (0.1% Triton X-100 in PBS) three times for 5 min each, they were incubated with secondary antibodies for 1 h at RT. Subsequently, the samples were washed in PBST three times for 5 min each, followed by staining DNA in 0.1 µg/ml DAPI (Roche) in PBS for 10 min at RT. After washing with PBS once, cells were mounted with VECTASHIELD Mounting Medium (Vector Laboratories).

For the analysis of K-fiber, RPE-1 cells were seeded onto 35 mm glass-bottom culture dishes (MatTek) for 24 to 36 h, and supplemented with MG132 (50 µM) for 1 h. After rinsing the cells with culture medium, the cells were incubated with CaCl_2_ buffer (1 mM MgCl_2_, 1 mM CaCl_2_, 0.5% Triton X-100, 100 mM Pipes, adjusted to pH 6.8) for 1 min at 37°C. The cells were then fixed in 1% glutaraldehyde (Nacalai Tesque) in PHEM for 10 min at 37°C. To quench the reaction, 0.1 g/ml sodium tetrahydridoborate (Nacalai Tesque) in PHEM was added and incubated for 20 min at RT. The cells were permeabilized with 0.5% Triton X-100 in PBS for 10 min and blocked with Abdil for 5 to 10 min at RT. The cells were incubated with primary antibody for 1 h at 37°C, followed by the aforementioned protocol for secondary antibody staining and counterstaining with DAPI.

For α-tubulin staining in error correction assay, RPE-1 or HeLa cells were seeded onto 35 mm glass-bottom culture dishes (MatTek) for 24 to 36 h. The cells were then fixed and permeabilized with 3.2% PFA, 0.5% Triton X-100, 1% glutaraldehyde in PHEM for 10 min at RT, followed by blocking with Abdil for 5 to 10 min at RT. Subsequently, the cells were incubated with primary antibodies for 1 h at 37°C. The secondary antibody staining and counterstaining with DAPI were performed as above.

For micronuclei analysis, MEFs or RPE-1 cells were seeded onto 35 mm glass-bottom culture dishes (MatTek) and cultured for two days. The cells were then fixed and permeabilized with a solution containing 4% PFA and 0.5% Triton X-100 in PHEM buffer for 10 minutes at RT. Nuclei were stained with DAPI as above.

For the analysis of Aurora B, H3T3ph, and H2AT120ph, chromosome spread samples were used. The cells were cultured with 100 ng/ml nocodazole (Sigma) for 2 to 4 h at 37°C. To spread and swell chromosomes, the method was modified from the protocol described previously (Erin C. Moran, 2021). After shaking off and collecting the mitotic-arrested cells, they were suspended in a hypotonic buffer (75 mM KCl (Nacalai Tesque): 0.8% sodium citrate (Nacalai Tesque): water at 1:1:1) with 1x cOmplete EDTA-free proteinase inhibitor (Roche) on ice. The suspended cells were cytospun onto coverslips using the Cytospin III Cytocentrifuge and fixed in 2% PFA in PBS for 20 min. The cells were blocked in 1% BSA in PBS for 10 min at RT, followed by incubation with primary antibody for 1 h at 37°C or overnight at 4°C. Subsequently, the above protocol for secondary antibody staining and counterstaining with DAPI was performed.

Primary antibodies used were diluted in Abdil, except for Aurora B (diluted in 1% BSA/PBS). The primary antibodies included rabbit anti-human Hec1 (1:5000) (Abcam, ab3613), mouse anti-human CENP-A (1:250) (Ando et al., 2002), mouse anti-human Aurora B (1:500) (BD Bioscience, 611082), rabbit anti-human DSN1 (1:2000) (a gift from Iain Cheeseman, Whitehead Institute, MIT) (Kline et al., 2006), rabbit anti-human KNL1 (1:2000) (a gift from Iain Cheeseman, Whitehead Institute, MIT) (Cheeseman et al., 2008), mouse anti-human Bub1 (1:400) (MAB3610, Millipore), rabbit anti-H2AT120ph (1:1000) (ActiveMotif), mouse anti-H3T3ph (1:3000) (Kimura et al., 2008), FITC-conjugated mouse anti-α-tubulin (1:1000) (Sigma, F2168), and mouse anti-α-tubulin (1:5000) (Sigma). Secondary antibodies used in immunofluorescence staining were FITC-conjugated goat anti-mouse IgG (1:1000) (Jackson ImmunoResearch), FITC-conjugated goat anti-rabbit IgG (1:1000) (Jackson ImmunoResearch), Alexa647-conjugated goat anti-mouse IgG (1:1000) (Jackson ImmunoResearch), and Alexa647-conjugated goat anti-rabbit IgG (1:1000) (Jackson ImmunoResearch).

Immunofluorescence images were captured with a spinning disk confocal unit CSU-W1 or CSU-W1-SoRa (Yokogawa) controlled with NIS-elements (v5.42.01, Nikon) with an objective lens (Nikon; PlanApo VC 60×/1.40 or Lambda 100×/1.45 NA) and an Orca-Fusion BT (Hamamatsu Photonics) sCMOS camera. The images were acquired with Z-stacks at intervals of 0.2 or 0.3-µm. Maximum intensity projections (MIPs) of the Z-stack were generated using Fiji software (Schindelin et al., 2012) for display and analysis. These images were processed using Fiji and Photoshop 2019 (Adobe).

### Live cell imaging

For analyzing CENP-C^WT^ or CENP-C^ΔM12BD^ RPE-1 cells, cells plated onto 35 mm glass-bottom dishes (MatTek) were switched to phenol red-free culture medium (phenol red-free DMEM (Nakalai Tesque), supplemented with 20% fetal bovine serum, 25 mM HEPES, 2 mM L-glutamine) and sealed with mineral oil (Sigma). Images were captured every 2 min with DeltaVision Elite imaging system (GE healthcare) equipped with a PlanApo N OSC 60×/1.40 NA oil immersion objective lens (Olympus) and a CoolSNAP HQ2 CCD camera (Photometrics) controlled with built-in SoftWoRx software (version 5.5) in a temperature-controlled room at 37°C. A Z-series of 7 sections with 2-µm increments was acquired. The Z-series was then projected using MIP for analysis by SoftWoRx. For analyzing CENP-T^WT^, CENP-T^ΔNBD-1^, or CENP-T^ΔNBD-2^ RPE-1 cells, the cells plated onto 35 mm glass-bottom dishes supplemented with IAA (500 µM) for two days were switched to phenol red-free culture medium supplemented with IAA (500 µM) and SPY505-DNA (1:1000) (Spirochrome) for 4 to 6 h before imaging. For analyzing *Cenpc*^+/+^, *Cenpc^+/ΔM12BD^*, or *Cenpc^ΔM12BD/ΔM12BD^*MEFs, the cells plated onto 35 mm glass-bottom dishes were switched to phenol red-free culture medium supplemented with SPY650-DNA (1:3000) (Spirochrome) for 4 to 6 h before imaging. Images were captured every 2 min with CSU-W1-SoRa system described above at 37°C under 5% CO_2_ condition. Z-series of 7 sections were taken at intervals of 2-µm. Acquired time-lapse images were projected by MIP and processed using Fiji and Photoshop 2019 (Adobe).

For chromosome oscillation analysis, cells were treated with SiRTubulin (1:5000) (Spirochrome) for 4 to 6 h and then supplemented with MG132 (5 µM) for 2 h. Images were filmed every 3 sec for 5 min, Z-series of 5 or 7 sections in 0.5-µm increments. The deviation from the average position (DAP) was determined according to the previously described method (Iemura and Tanaka, 2015; Stumpff et al., 2008). In brief, the Z-stack images were deconvolved using plug-in function of NIS-element (Richardson-Lucy method) and then projected by MIP for quantification. Individual kinetochores were tracked by Manual Tracking (plug-in of Fiji, http://rsb.info.nih.gov/ij/plugins/track/track.html) after aligning the cell movement using the StackReg (plug-in of Fiji) (Thevenaz et al., 1998). The obtained data were analyzed in Microsoft Excel.

### Flow cytometry

MEFs were collected after trypsinization. The harvested cells were washed twice with ice-cold 1% BSA/PBS, fixed with ice-cold 70% ethanol, and stored at -20°C. The fixed cells were washed again with 1% BSA/PBS and incubated with 20 μg/ml propidium iodide (Sigma) in 1% BSA/PBS for 20 min at RT, followed by overnight incubation at 4°C. The stained cells were then subjected to flow cytometry analysis using a BD FACS Canto II Flow Cytometer and analyzed with BD FACSDiva 9.0 Software (BD Biosciences).

### Cell viability assay

RPE-1 cells were seeded onto opaque-walled tissue culture plates with a clear bottom (Corning) at a density of 2500 cells per well and incubated for 12 h. The cells were treated with nocodazole at indicated concentrations for 3 days, with each concentration tested in triplicate. After treatment, RealTime-Glo reagents (RealTime-Glo MT Cell Viability Assay, Promega) were added to the cells and incubated for 1 h at 37°C. Luminescence was measured using a GloMax Discover System Microplate Reader (Promega). IC_50_ values were determined using GraphPad Prism 9.5.1 (GraphPad).

### Error correction assay

RPE-1 or HeLa cells were plated and incubated on 35 mm glass-bottom culture dishes (MatTek) for 24 to 36 h. Subsequently, the cells were treated with 50 µM monastrol (Selleckchem) for 4 h, followed by washing with culture medium four times. The cells were then supplemented with MG132 (5 µM) (Sigma) for indicated timing and fixed (3.2% PFA, 0.5% Triton X-100, 1% glutaraldehyde in PHEM at RT for 10 min) immediately. Following fixation and permeabilization, staining was performed using antibodies and DAPI as aforementioned protocol.

### Quantification and Statistical Analysis

The fluorescence signal intensities of DSN1, KNL1, Hec1, Bub1, and mScarlet-DSN1 on kinetochores were quantified using Imaris (Bitplane). The fluorescence signal intensities of Aurora B and H3T3ph within the inner centromere regions were measured using Fiji. The signal intensities were quantified by subtracting background signals in a 15-pixel circular region of chromosome adjacent to the inner centromere region from the signals in a 15-pixel circular region of the inner centromere region. The fluorescence signal intensities of H2AT120ph within the kinetochore-proximal centromere region were measured using Fiji. The signal intensities were quantified by subtracting background signals in a chromosomal region from the signals in a 10-pixel circular region circular area centered on the kinetochore. For quantification of the k-fiber signal intensities, to select the spindle area in a cell thresholding is applied to the tubulin signals (MIP) using Fiji. The mean signal intensities in the selected area are measured. The value is subtracted by the mean background signal intensities adjacent to the spindle area and quantified as the averaged k-fiber signals. Data processing was carried out using Microsoft Excel and GraphPad Prism 9.5.1 (GraphPad), and p-values were calculated using two-tailed Student’s t-test or one-way or two-way ANOVA test followed by multiple comparison tests. Each experiment was repeated four times (Figure 4F), three times (Figures 3C, 3D, 3I, 3J, 4B, 4E, 5A, 5B, 5D, 5F, 7B), or twice (Figures 3B, 4C, 4D, 5E, 7C, 7D) when not mentioned in figure legends, and representative data of replicates were presented.

## Supplementary Information

The article contains 6 Supplementary Figures and 1 Supplementary Table.

## Acknowledgments

The authors are very grateful to members of the Fukagawa Lab for the fruitful discussion. We also thank R. Fukuoka, K. Oshimo, Y. Kubota, M. Nakagawa, A. Kanie, N. Yasue, and S. Oka for their technical assistance. This work was supported by CREST of JST (JPMJCR21E6), JSPS KAKENHI Grant Numbers 20H05389, 21H05752, 22H00408, 22H04692, and 24H02281 to TF, JSPS KAKENHI Grant Numbers 16K18491, 21H02461 22H04672, and 24K02005, Takeda Science Foundation and Daiichi Sankyo Foundation of Life Science to MH. This work was also supported by JSPS KAKENHI Grant Number JP22H04926 Grant-in-Aid for Transformative Research Areas ― Platforms for Advanced Technologies and Research Resources “Advanced Bioimaging Support” and the NIBB Collaborative Research Program (19-352, 20-305, 21-238, 22NIBB327). The images of mouse and pipette in Figure 2A are from TogoTV (© 2016 DBCLS TogoTV, CC-BY-4.0).

## Author contributions

Conceptualization: M.Hara., T.Fukagawa; Investigation: W.K., M.Hara., K.O., Y.Tokunaga, Y.H., J.M. ; Formal analysis: W.K., M.Hara, Y.H., K.O. Y.Tokunaga; Resources: Y.Takenoshita., M.Hashimoto., H.S., T.Fujimori, Y.W; Data curation: W.K., M.Hara; Writing - original draft: M. Hara, T., Fukagawa; Writing - review & editing: M.Hara, T. Fukagawa; Supervision: M. Hara T.Fukagawa; Project administration: T.Fukagawa; Funding acquisition: M. Hara, T.Fukagawa

## Declaration of Interests

All authors declare no competing interests.

## Legends for Supplementary Figures

**Supplementary Figure 1.**
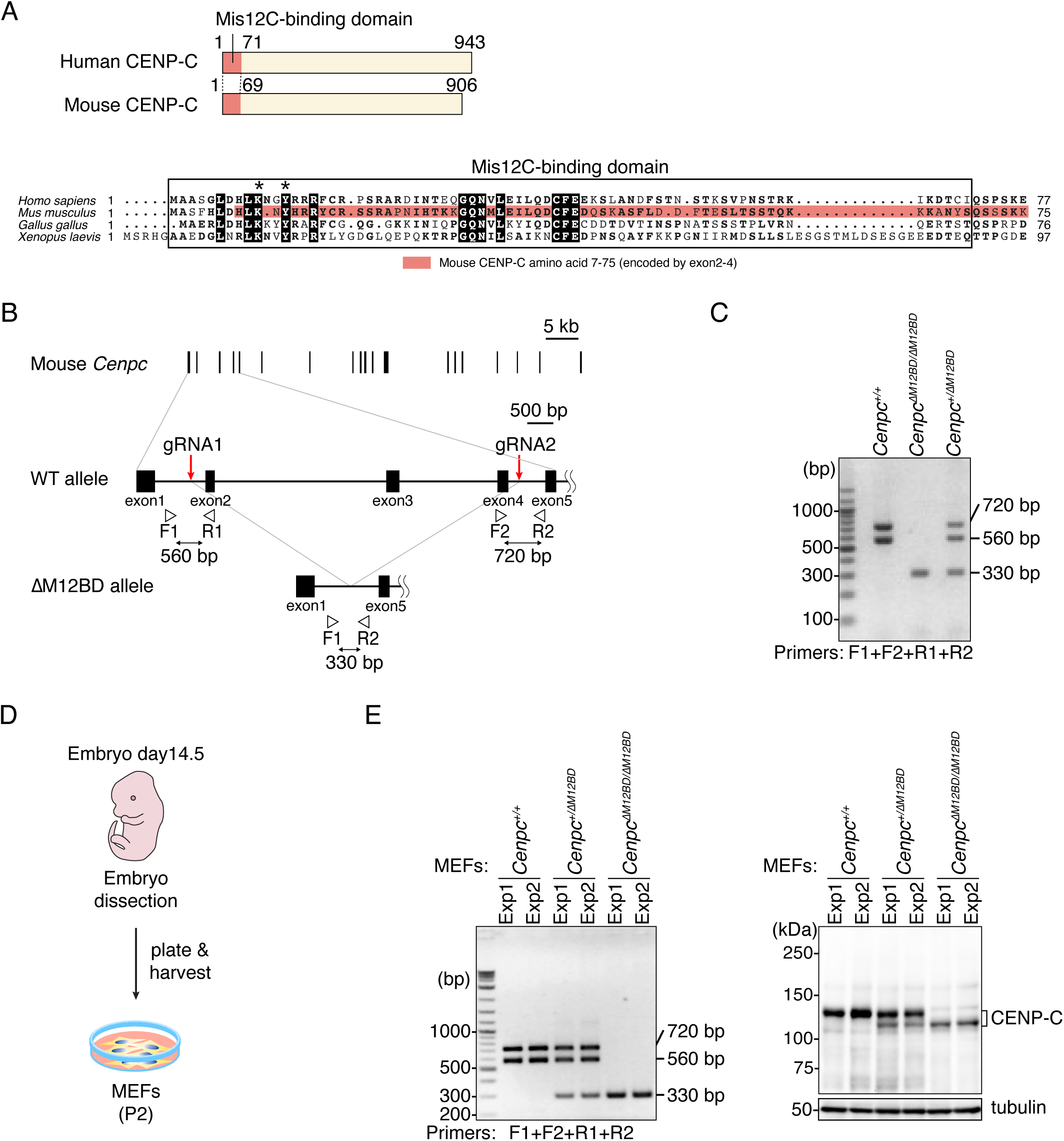
Generation of CENP-C mutant mice and MEFs. (A) Alignment of the amino acid sequence of the CENP-C N-terminus including Mis12C-binding domain. The Mis12C-binding domain of human CENP-C and the corresponding region in CENP-C of other species are boxed. Asterisks show the conserved amino acids in CENP-C essential for Mis12C binding (lysine 10 and tyrosine 13) (Screpanti et al., 2011). In mouse CENP-C, most of the Mis12C-binding domain, including the conserved amino acid for Mis12C-binding, are encoded in exon2-4 (aa 7-75). *Homo sapience* CENP-C: NP_001803, *Mus musculus* CENP-C: NP_031709, *Gallus gallus* CENP-C: NP_001376225.2, *Xenopus laevis* CENP-C: NP_0011594. (B) Schematic representation of the exon 2-4 deletion from mouse *Cenpc* gene. mouse *Cenpc1* (*Cenpc*) gene has 19 exons. Using the CRISPR-Cas9 system with two gRNAs, the exon2-4 is deleted to make the Mis12C-binding domain deletion mutant (ΔM12BD). The position of gRNAs and PCR primers for genotyping PCR are shown. The amplicon sizes are approximately 560 bp (F1/R1), 720 bp (F2/R2), 330 bp (F1/R2). (C) Genotyping PCR of *ΔM12BD Cenpc* mutant mice (*Cenpc^ΔM12BD^*). (D) Schematic representation of the establishment of MEFs from embryos. (E) Genotyping PCR and immunoblotting for CENP-C of *Cenpc^ΔM12BD^* MEFs. The MEFs isolated from two independent experiments were tested. In immunoblots, mouse CENP-C was detected by an antibody against mouse CENP-C, and α-tubulin was probed as a loading control.

**Supplementary Figure 2.**
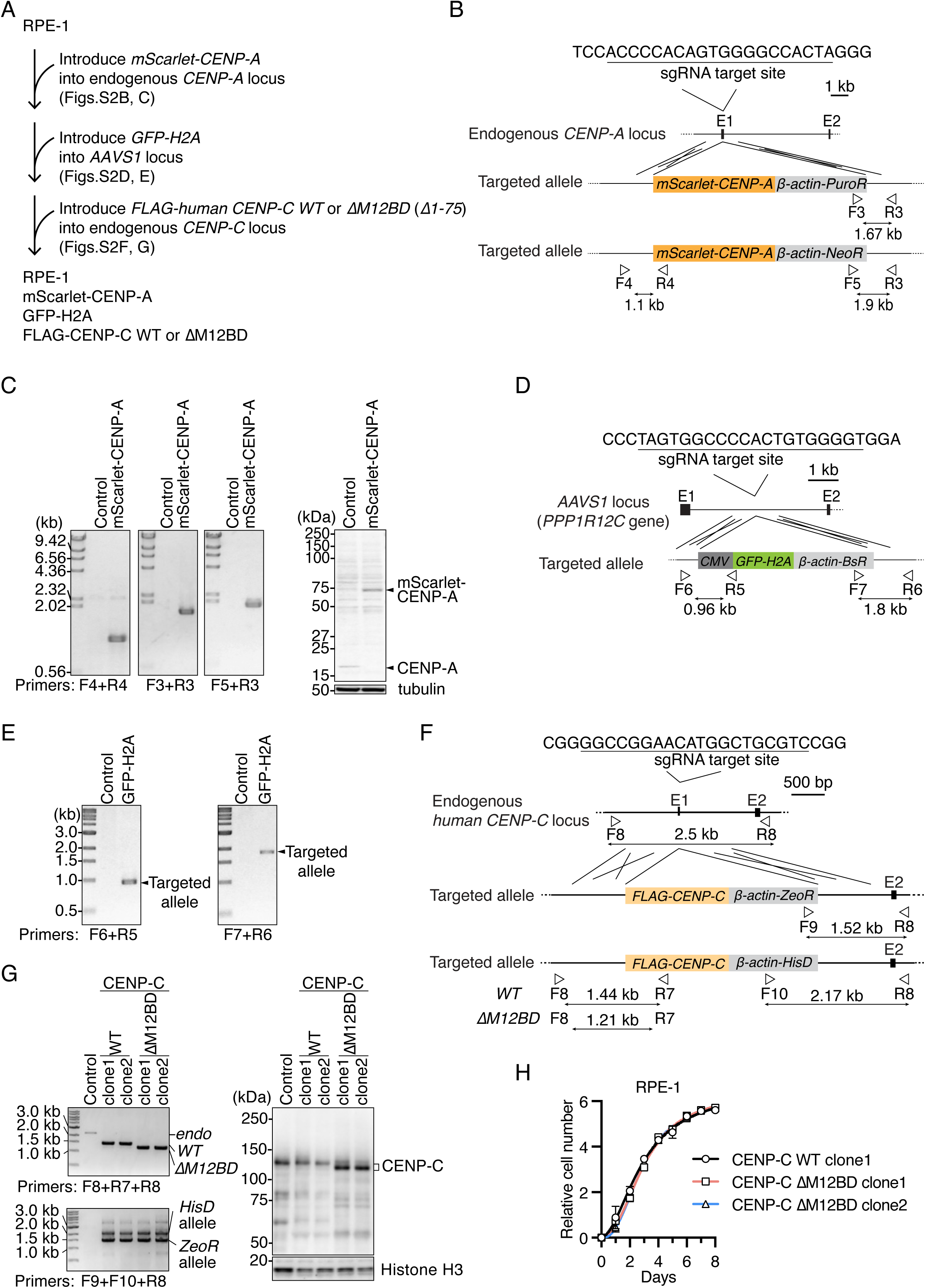
Generation of CENP-C mutant RPE1 cell lines. (A) Strategy to generate CENP-C^WT^ or CENP-C^ΔM12BD^ RPE1 cells expressing mScarlet-CENP-A and GFP-H2A (See also Figure 3). (B) Schematic representation of *mScarlet-CENP-A* cDNA targeting into the endogenous *CENP-A* locus. To express mScarlet-fused CENP-A under the control of the endogenous *CENP-A* promoter, *mScarlet-CENP-A* cDNA was targeted into the exon1 by CRISPR/Cas9-mediated homologous recombination. Since the targeting constructs have puromycin resistance genes (*PuroR*) or neomycin resistance genes (*NeoR*), targeted cells were selected using these selection markers. The gRNA sequence and position of primers for genotyping are shown. (C) Genotyping PCR and immunoblotting for mScarlet-CENP-A in *mScarlet-CENP-A*-introduced RPE-1 cells. The genotype in isolated single clones was examined using the primers shown in (B). In immunoblots, CENP-A was detected by an antibody against CENP-A, and α-tubulin was probed as a loading control. (D) Schematic representation of *GFP-H2A* cDNA targeting into the *AAVS1* locus (*PPP1R12C* gene). To express GFP-fused histone H2A from the *AAVS1* locus, *GFP-H2A* cDNA was targeted into the intron1 by CRISPR/Cas9-mediated homologous recombination. Since the targeting construct has blasticidin resistance genes (*BsR*), targeted cells were selected using the *BsR* marker. The gRNA sequence and position of primers for genotyping are shown. (E) Genotyping PCR of *GFP-H2A*-introduced RPE-1 cells. The genotype in an isolated single clone was examined using the primers shown in (D). (F) Schematic representation of *FLAG-human CENP-C* cDNA targeting into the endogenous *human CENP-C* locus. To express FLAG-tagged CENP-C wiled-type (WT) or a Mis12C-binding domain deletion mutant (ΔM12BD: Δ1-75) under the control of the endogenous *CENP-C* promoter, *FLAG-CENP-C WT* or *ΔM12BD* cDNA was targeted into the exon1 by CRISPR/Cas9-mediated homologous recombination. Since the targeting constructs have Zeocin resistance genes (*ZeoR*) or histidinol resistance genes (*histidinol dehydrogenase*: *HisD*), targeted cells were selected using these selection markers. The gRNA sequence and position of primers for genotyping are shown. (G) Genotyping PCR and immunoblotting for FLAG-CENP-C in *FLAG-CENP-C*-introduced RPE-1 cells. The genotype in isolated single clones was examined using the primers shown in (F). In immunoblots, CENP-C was detected by an antibody against human CENP-C, and histone H3 was probed as a loading control. (H) The growth curve of CENP-C^WT^ or CENP-C^ΔM12BD^ RPE1 cells. The cell numbers were normalized to those at Day 0 of each line. Error bars indicate the mean and standard deviation. Two clones of CENP-C^ΔM12BD^ RPE1 cells were examined.

**Supplementary Figure 3.**
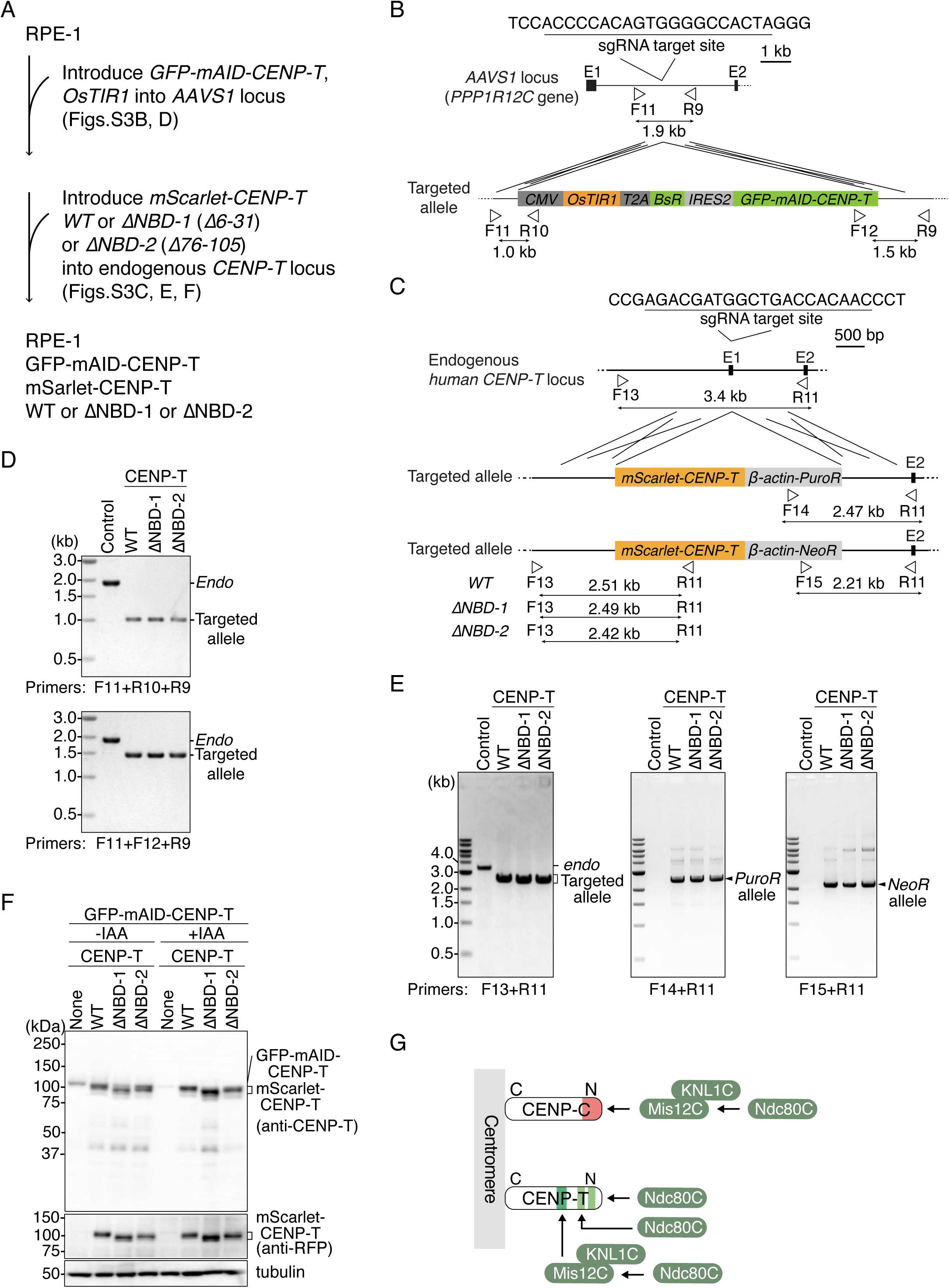
Generation of CENP-T mutant RPE1 cell lines. (A) Strategy to generate CENP-T^WT^, CENP-T^ΔNBD-1^, or CENP-T^ΔNBD-2^ RPE1 cells (See also Figure 4). In the cells, mini auxin-inducible degron (mAID)-tagged human CENP-T was expressed together with OsTIR1. Since mAID-fused CENP-T is degraded upon IAA (Indole-3-acetic Acid) treatment, mScarlet-tagged CENP-T is only expressed CENP-T in CENP-T^WT^, CENP-T^ΔNBD-1^, or CENP-T^ΔNBD-2^ RPE1 cells. (B) Schematic representation of *GFP-mAID-CENP-T* and *OsTIR1* cDNA targeting into the *AAVS1* locus (*PPP1R12C* gene). To express GFP and mAID-fused human CENP-T and OsTIR1, the expression cassette was targeted into the intron1 by CRISPR/Cas9-mediated homologous recombination. Since the targeting construct has blasticidin resistance genes (*BsR*), targeted cells were selected using the *BsR* marker. The gRNA sequence and position of primers for genotyping are shown. (C) Schematic representation of *mScarlet-CENP-T* cDNA targeting the endogenous *CENP-T* locus. To express mScarlet-tagged CENP-T wild-type (WT) or each of Ndc80C-binding domain mutants (ΔNBD-1: Δ6-31, ΔNBD-2: Δ76-105) under the control of the endogenous *CENP-T* promoter, *mScarlet-CENP-T WT*, *ΔNBD-1* or *ΔNBD-2* cDNA was targeted into the exon1 by CRISPR/Cas9-mediated homologous recombination. Since the targeting constructs have puromycin resistance genes (*PuroR*) or neomycin resistance genes (*NeoR*), targeted cells were selected using these selection markers. The gRNA sequence and position of primers for genotyping are shown. (D) Genotyping PCR for *GFP-mAID-CENP-T/OsTIR1* expression cassette in CENP-T^WT^, CENP-T^ΔNBD-1^, or CENP-T^ΔNBD-2^ RPE1 cells. The genotype in isolated single clones was examined using the primers shown in (B). Wild-type RPE-1 cells were used as a control. (E) Genotyping PCR for targeted *mScarlet-CENP-T* in CENP-T^WT^, CENP-T^ΔNBD-1^, or CENP-T^ΔNBD-2^ RPE1 cells. The genotype in isolated single clones was examined using the primers shown in (C). Wild-type RPE-1 cells were used as a control. (F) CENP-T protein expression in CENP-T^WT^, CENP-T^ΔNBD-1^, or CENP-T^ΔNBD-2^ RPE-1 cells. The cells were treated with or without IAA for one day and examined GFP-mAID-CENP-T and mScarlet-CENP-T protein expression using an antibody against CENP-T or mScarlet (RFP). α-tubulin was probed as a loading control. (G) Schematic representation of recruitment of KMN subcomplexes onto CENP-C or CENP-T in the human cells.

**Supplementary Figure 4.**
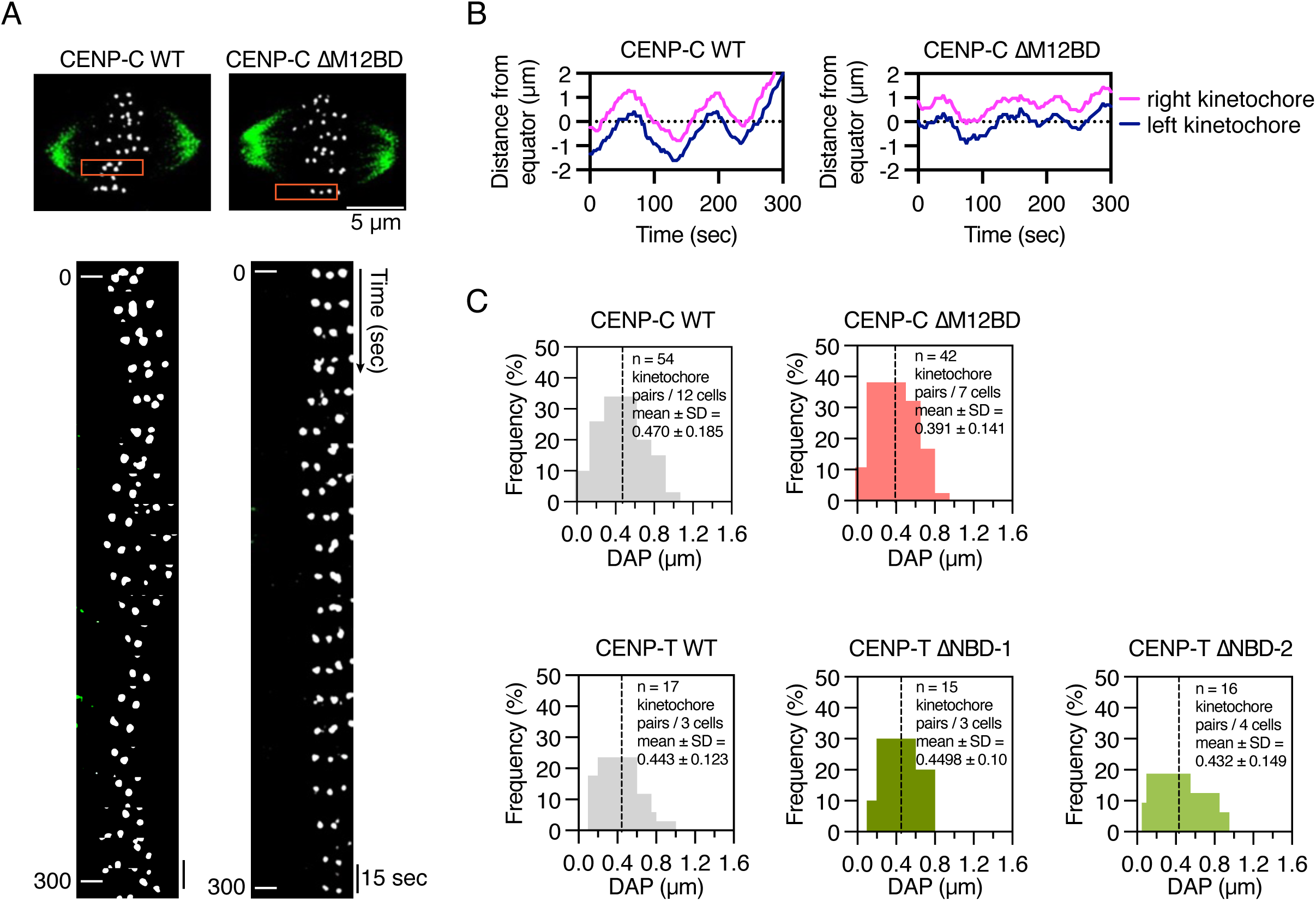
Chromosome oscillation is dampened in CENP-C^ΔM12BD^ RPE1 cells. (A) Chromosome oscillation in CENP-C^WT^ or CENP-C^ΔM12BD^ cells. Representative cell images are shown. mScarlet-CENP-A is a kinetochore marker (gray). The spindle was visualized with SiR-Tubulin (green). Scale bar, 5 μm. Kymographs represent the boxed kinetochore pairs at 15 sec intervals. (B) Distance of sister kinetochores from the equator in CENP-C^WT^ or CENP-C^ΔM12BD^ cells. Representative trajectories for kinetochore pairs are shown. (C) Frequency of DAP (the deviation from average position) values in CENP-C^WT^ or CENP-C^ΔM12BD^ cells. DAP is calculated by using trajectories of kinetochore pairs. 54 kinetochore pairs in 12 CENP-C^WT^ cells and 42 kinetochore pairs in 7 CENP-C^ΔM12BD^ cells were used for the quantification. The dashed lines indicate the mean. (D) Frequency of DAP values in CENP-T^WT^, CENP-T^ΔNBD-1^, or CENP-T^ΔNBD-2^ cells. 17 kinetochore pairs in 3 CENP-T^WT^ cells, 15 kinetochore pairs in 3 CENP-T^ΔNBD-1^ cells, and 16 kinetochore pairs in 4 CENP-T^ΔNBD-2^ cells were used for the quantification. The dashed lines indicate the mean.

**Supplementary Figure 5.**
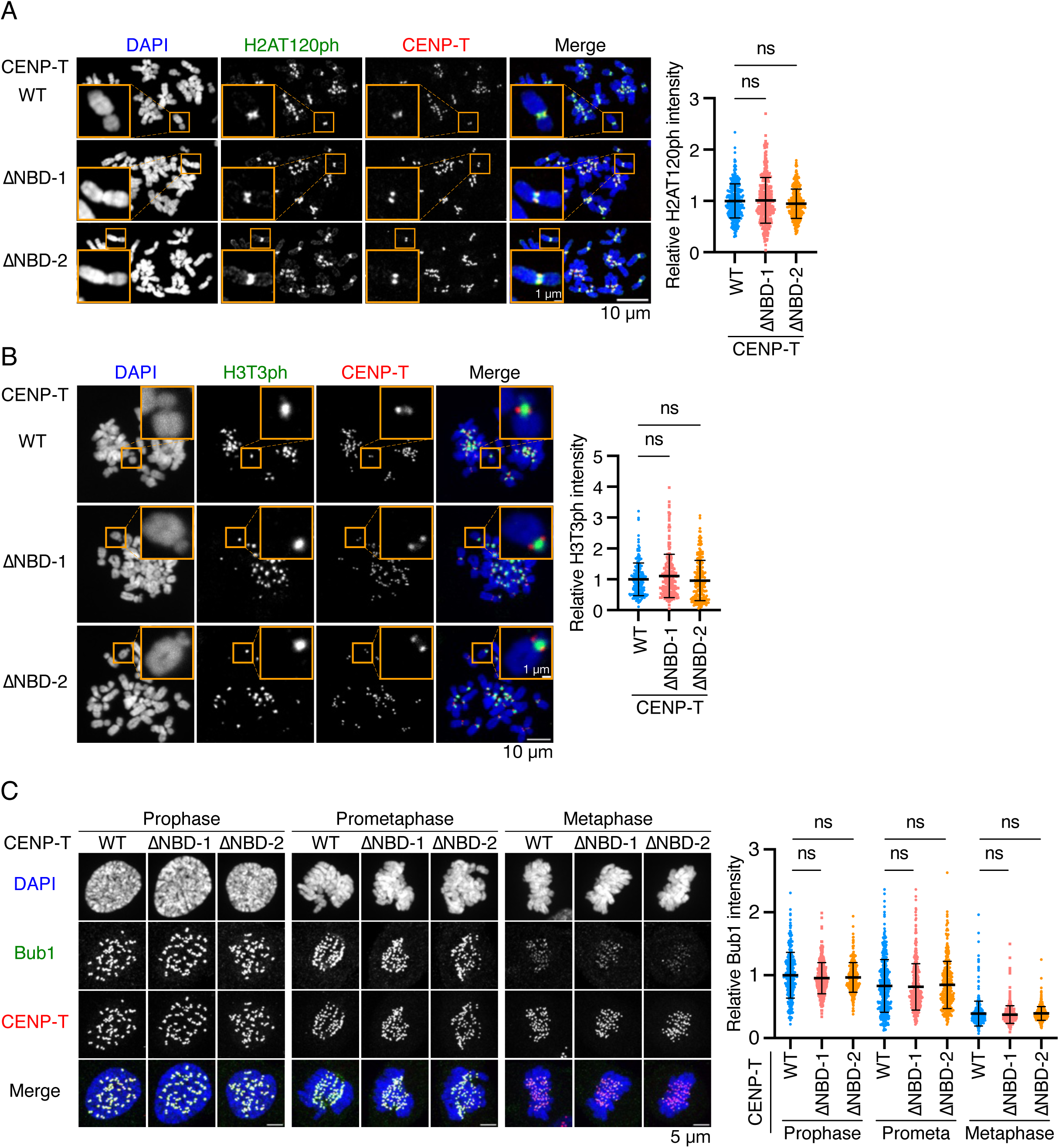
Bub1, H2At120ph, and H3T3ph localization in CENP-T mutant RPE1 cell lines. (A) H2AT120ph localization in CENP-T^WT^, CENP-T^ΔNBD-1^, or CENP-T^ΔNBD-2^ RPE-1 cells. H2AT120ph was stained with an antibody against H2AT120ph (green). mScarlet-CENP-T is a kinetochore marker (CENP-T, red). DNA was stained with DAPI (blue). Scale bar, 10 μm. The insets show an enlarged single chromosome (Scale bar, 1 μm). H2AT120ph signal intensities at kinetochore-proximal centromeres were quantified. A representative result from three independent experiments is shown (Mean and SD, one-way ANOVA with Dunnett’s multiple comparison test, CENP-T^WT^: n = 316 kinetochores from 5 cells, CENP-T^ΔNBD-1^: n = 306 kinetochores from 5 cells, CENP-T^ΔNBD-2^: n = 325 kinetochores from 5 cells). (B) H3T3ph localization in CENP-T^WT^, CENP-T^ΔNBD-1^, or CENP-T^ΔNBD-2^ RPE1 cells. H3T3ph was stained with an antibody against H3T3ph. H3T3ph localization at centromeres was examined and quantified as in (A). Scale bar, 10 μm. A representative result from two independent experiments is shown (Mean and SD, one-way ANOVA with Dunnett’s multiple comparison test, CENP-T^WT^: n = 195 centromeres from 5 cells, CENP-T^ΔNBD-1^: n = 206 centromeres from 5 cells, CENP-T^ΔNBD-2^: n = 210 centromeres from 5 cells). (C) Bub1 localization in CENP-T^WT^, CENP-T^ΔNBD-1^, or CENP-T^ΔNBD-2^ RPE1 cells. Bub1 was stained with an antibody against Bub1 (green). mScarlet-CENP-T is a kinetochore marker (CENP-T, red). DNA was stained with DAPI (blue). Scale bar, 5 μm. Bub1 signal intensities at kinetochores were quantified. A representative result from two independent experiments is shown (Mean and SD, one-way ANOVA with Dunnett’s multiple comparison test, CENP-T^WT^: n = 427 kinetochores from 5 cells, CENP-T^ΔNBD-1^: n = 424 kinetochores from 5 cells, CENP-T^ΔNBD-2^: n = 423 kinetochores from 5 cells).

**Supplementary Figure 6.**
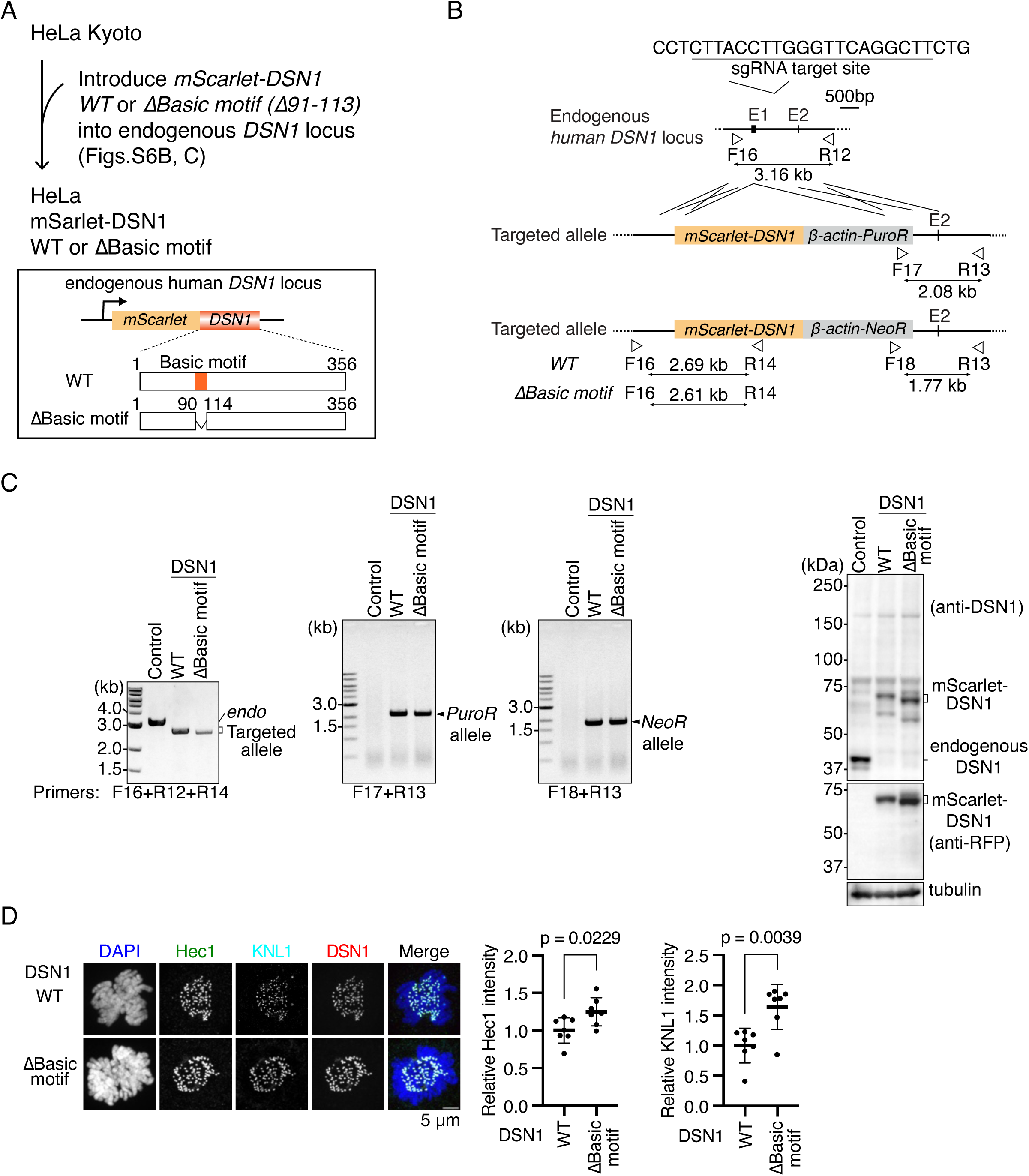
Generation of DSN1 mutant HeLa cell lines. (A) Strategy to generate DSN1^WT^ or DSN1^ΔBasic^ ^motif^ HeLa cells. (See also Figure 7A). (B) Schematic representation of *mScarlet-DSN1* cDNA targeting the endogenous *DSN1* locus. To express mScarlet-tagged DSN1 wiled-type (WT) or a basic motif deletion mutant (ΔBasic motif: Δ91-113) under the control of the endogenous *DSN1* promoter, *mScarlet-DSN1 WT* or *ΔBasic motif,* cDNA was targeted into *DSN1 locus* by CRISPR/Cas9-mediated homologous recombination. Since the targeting constructs have puromycin resistance genes (*PuroR*) or neomycin resistance genes (*NeoR*), targeted cells were selected using these selection markers. The gRNA sequence and position of primers for genotyping are shown. (C) Genotyping PCR and immunoblotting for mScarlet-DSN1 in *mScarlet-DSN1*-introduced HeLa cells. The genotype in isolated single clones was examined using the primers shown in (B). In immunoblots, DSN1 was detected by an antibody against DSN1 or mScarlet (RFP), and α-tubulin was probed as a loading control. (D) Hec1 and KNL1 localization in DSN1^WT^ or DSN1^ΔBasic^ ^motif^ HeLa cells. Hec1, a subunit of the Ndc80C (green), and KNL1, a subunit of the KNL1C (cyan), were stained with antibodies against Hec1 and KNL1, respectively. mScarlet-DSN1 is a kinetochore marker (DSN1, red). DNA was stained with DAPI (blue). Scale bar, 5 μm. Hec1 or KNL1 signal intensities at mitotic kinetochores were quantified. A representative result from two independent experiments is shown (Mean and SD, two-tailed Student’s t-test, DSN1^WT^: n = 7, DSN1^ΔBasic^ ^motif^: n = 7).

**Table S1.**
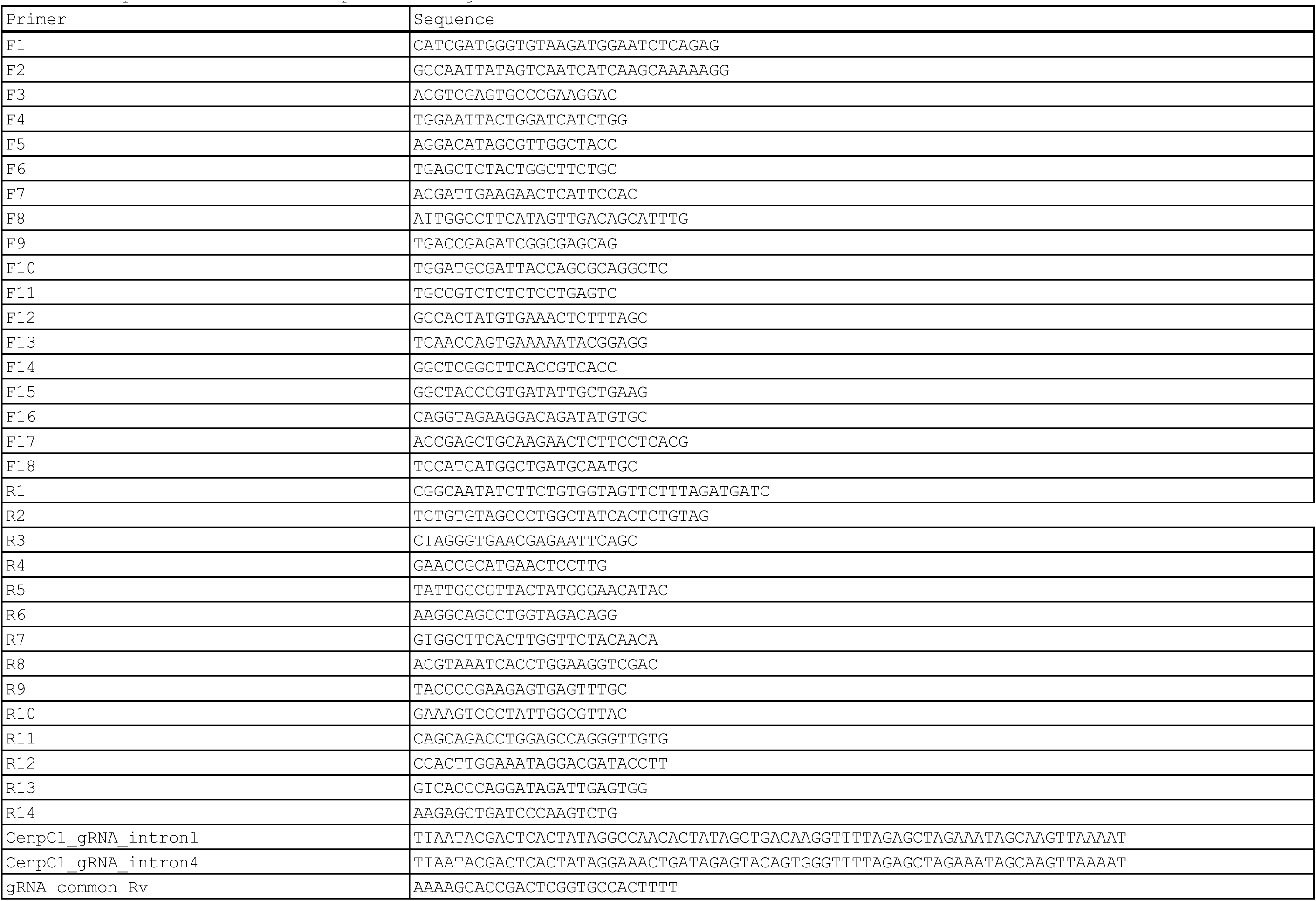
Sequence information of primers and gRNAs.

## Notes

### Competing Interest Statement

The authors have declared no competing interest.

